# Immunologic perturbations in severe COVID-19/SARS-CoV-2 infection

**DOI:** 10.1101/2020.05.18.101717

**Authors:** Leticia Kuri-Cervantes, M. Betina Pampena, Wenzhao Meng, Aaron M. Rosenfeld, Caroline A.G. Ittner, Ariel R. Weisman, Roseline Agyekum, Divij Mathew, Amy E. Baxter, Laura Vella, Oliva Kuthuru, Sokratis Apostolidis, Luanne Bershaw, Jeannete Dougherty, Allison R. Greenplate, Ajinkya Pattekar, Justin Kim, Nicholas Han, Sigrid Gouma, Madison E. Weirick, Claudia P. Arevalo, Marcus J. Bolton, Eileen C. Goodwin, Elizabeth M. Anderson, Scott E. Hensley, Tiffanie K. Jones, Nilam S. Mangalmurti, Eline T. Luning Prak, E. John Wherry, Nuala J. Meyer, Michael R. Betts

## Abstract

Although critical illness has been associated with SARS-CoV-2-induced hyperinflammation, the immune correlates of severe COVID-19 remain unclear. Here, we comprehensively analyzed peripheral blood immune perturbations in 42 SARS-CoV-2 infected and recovered individuals. We identified broad changes in neutrophils, NK cells, and monocytes during severe COVID-19, suggesting excessive mobilization of innate lineages. We found marked activation within T and B cells, highly oligoclonal B cell populations, profound plasmablast expansion, and SARS-CoV-2-specific antibodies in many, but not all, severe COVID-19 cases. Despite this heterogeneity, we found selective clustering of severe COVID-19 cases through unbiased analysis of the aggregated immunological phenotypes. Our findings demonstrate broad immune perturbations spanning both innate and adaptive leukocytes that distinguish dysregulated host responses in severe SARS-CoV-2 infection and warrant therapeutic investigation.

**One Sentence Summary:** Broad immune perturbations in severe COVID-19

## Introduction

The coronavirus-19-disease (COVID-19) pandemic caused by the severe acute respiratory syndrome coronavirus 2 (SARS-CoV-2) has surpassed four million cases world-wide (4,088,842 as of 05/12/2020), causing more than 283,000 deaths in 215 countries (*1*). While asymptomatic in some, SARS-CoV-2 infection can cause viral pneumonia that progresses to acute respiratory distress syndrome (ARDS), and even multi-organ failure, in severe cases (*2, 3*). Reports have shown that SARS-CoV-2 has the ability to productively infect lung epithelium, gut enterocytes and endothelium (*4–6*). It is unclear whether disease severity is caused by the viral infection, the host response, or both, emphasizing the urgent need to understand the immune perturbations induced by SARS-CoV-2 (*3*). Knowledge of the immunological signatures of severe COVID-19 is continually evolving. Although lymphopenia has been linked to disease severity, the majority of published studies are based on retrospective analyses of clinical data (*3, 7–14*).

Immune profiling studies to date have been conducted as single case reports or focused only on moderate, severe or recovered COVID-19 with limited numbers of individuals (*15–18*), and have not necessarily reflected the range of comorbidities globally associated with severe COVID-19. Studies of peripheral blood mononuclear cells by mass cytometry or single cell RNA sequencing (scRNAseq) have provided valuable insights into possible immune perturbations in COVID-19 but have not assessed the contributions of granulocytic populations, or, in the case of scRNAseq, defined expression or modulation of cellular proteins (*16*). In particular, modulation of granulocytic populations is suggested to be relevant during COVID-19 infection (*12*).

To address these issues, we conducted a comprehensive analysis of the overall immunologic state of 42 individuals with different trajectories of SARS-CoV-2 infection and COVID-19 (moderate, severe, and recovered), compared with 12 healthy donors using whole blood to capture the full breadth of immunological perturbations and activation occurring in circulating lymphocytes and major granulocyte populations. We further explored modulation of the B cell repertoire, its associations with the establishment of a SARS-CoV-2-specific humoral response, and activation of T cells relative to disease severity. Together our results reveal a potential platform for assessing disease trajectory, and identify distinct immune perturbation patterns in severe COVID-19 that merit consideration for therapeutic immunomodulation strategies to ameliorate disease severity and organ failure.

## RESULTS

### Demographics and clinical characteristics of moderate and severe COVID-19+ individuals

We recruited 35 inpatients with active COVID-19, seven of whom had moderate and 28 with severe disease, seven recovered COVID-19+ donors, and 12 healthy donors (HD). All recovered donors reported mild disease, and did not receive inpatient care or COVID-19 directed therapy during the course of their illness. For inpatients, median follow up after enrollment was 27 days (range 20 – 43) since blood draw. General demographics and clinical characteristics are shown in Table 1. The median ages in the moderate and severe COVID-19+ groups were 59 and 68 years old, respectively, concordant with previous reports (*8*), and were not significantly different (p=0.51). Both the HD and recovered groups were significantly younger than individuals with severe COVID-19+ (p<0.001 in both cases). In line with a recent publication (*9*), the majority of the individuals in the severe and recovered groups were male (67.9% and 71.4%, respectively), while approximately 29% were male in the moderate disease group. The median number of days since onset of symptoms to disease progression in donors with severe COVID-19 was nine, similar to previous publications (*3, 10*). Individuals with moderate disease also reported a median of nine days since onset of symptoms. In accordance with a recent report (*19*), individuals with COVID-19 had high incidence of underlying pulmonary disease (11/35 including moderate and severe, 31.4%) and were current or former smokers (13/35 including moderate and severe, 42.7%, higher in individuals who developed severe disease).

**Table 1.** Demographics and clinical characteristics.

| **Characteristic** | **HD** | **Recovered** | **Moderate** | **Severe<sup>a</sup>** |
| --- | --- | --- | --- | --- |
| **n** | 12 | 7 | 7 | 28 |
| **Age** | 36 (24-61) | 30 (20-49) | 59 (29-64) | 68 (38-81) |
| **Male** | 6 (46.1) | 5 (71.4) | 2 (28.6) | 19 (67.9) |
| **Race** |  |  |  |  |
| Black or African American | - | - | 5 (71.4) | 16 (57.1) |
| Asian or Asian American | - | - | 0 | 2 (7.1) |
| White or Caucasian | - | - | 2 (14.3) | 11 (39.2) |
| **Past smoking history** | - | - | 2 (14.3) | 13 (46.4) |
| **Comorbidity** |  |  |  |  |
| Obesity | - | - | 3 (42.9) | 8 (28.6) |
| Hypertension | - | - | 5 (71.4) | 21 (75) |
| Diabetes | - | - | 1 (14.3) | 7 (25) |
| Thromboembolic complications | - | - | 1 (14.3) | 7 (25) |
| Coronary artery disease/myocardial infarction | - | - | 0 | 3 (10.7) |
| Underlying lung disease | - | - | 4 (57.1) | 7 (25) |
| Renal insufficiency/chronic kidney disease | - | - | 2 (14.2) | 20 (71.4) |
| Hyperlipidemia | - | - | 2 (71.4) | 14 (50) |
| **Treatment** |  |  |  |  |
| Hydroxychloroquine | - | - | 4 (57.1) | 25 (89.3) |
| Remdesivir<sup>b</sup> | - | - | 1 (14.2) | 12 (42.9) |
| **Days since onset of symptoms<sup>d</sup>** | - | 27 (17-32) | 9 (1-16) | 9 (1-25) |
| **Oxygen therapy/ARDS** |  |  |  |  |
| Nasal cannula | - | - | 3 (42.9) | 0 |
| HFNC / NIV | - | - | - | 4 (4.3) |
| Ventilator non-ARDS | - | - | - | 2 (67.1) |
| Mild ARDS | - | - | - | 3 (10.7) |
| Moderate ARDS | - | - | - | 9 (32.1) |
| Severe ARDS | - | - | - | 10 (35.7) |
| ECMO | - | - | - | 1 (3.6) |
| **Mortality** | 0 | 0 | 0 | 4 (14.3) |

Data are shown as number and percentage, n (%). Age is reported in median years (min-max).
Days since onset of symptoms is reported as median (min-max). Not all data were collected for
HD and recovered individuals. ARDS, acute respiratory distress syndrome; HFNC - NIV, high flow nasal cannula - non-invasive; ECMO, extracorporeal membrane oxygenation.

Hypertension and hyperlipidemia were the most frequent co-morbidities in moderate and severe COVID-19. The majority of individuals with severe COVID-19 presented with moderate and severe ARDS (*20*), and hospital mortality was 14.3% within this group. Thromboembolic complications, metabolic, vascular and pulmonary disease were also observed more frequently among those with severe disease (Table 1). As part of clinical care, D-dimer, procalcitonin, ferritin, lactate dehydrogenase, and C-reactive protein levels were measured in moderate and severe COVID-19 individuals. Median levels of D-dimer at the time of blood draw were 3.985 µg/ml in severe, and 0.62 µg/ml in moderate COVID-19 donors (severe n=20, moderate n=5; p=0.0022). We found higher levels of ferritin in the severe group compared to the moderate group (medians: 919.5 ng/ml in severe, n=20, and 162 ng/ml in moderate, n=5; p=0.007). Consistent with previous findings (*13*), median procalcitonin values were relatively low, though higher in severe donors than in those with moderate disease (medians of 0.45 ng/ml, n=15, and 0.06 ng/ml, n=5, respectively; p=0.0014). Levels of lactate dehydrogenase and C-reactive protein were similar across groups. Bacterial co-infection was present in nine individuals with severe COVID-19, and in only one moderate donor. An extended list of clinical information of the analyzed individuals is shown in Table S1.

### Immune perturbation in severe COVID-19

To assess the general landscape of immune responses and their perturbation during severe COVID-19, we performed extensive immunophenotyping to characterize the frequencies of circulating immune subsets in HD, or in moderate, severe and recovered COVID-19 individuals (Fig. 1, Fig. S1). We observed an expansion in the proportion of both neutrophil and eosinophil populations in severe COVID-19 donors compared to HD (median neutrophil frequencies within viable CD45+ cells: 79.9% in severe COVID-19 and 47.7% in HD; p<0.0001; and, median eosinophil frequencies within viable CD45+ cells: 0.68% eosinophils in severe COVID-19 and 0.17% in HD, p=0.0015; Fig. 1A-C). The neutrophil frequency also differed significantly between moderate vs. severe COVID-19 disease (p=0.0046, median frequency of 53% of viable CD45+ in moderate group), but did not show increased activation or cycling (Fig. S2A). Furthermore, we saw decreased expression of CD15 in neutrophils between HD and severe COVID-19 individuals (p=0.0095), but not in eosinophils (Fig. S2B). We did not observe significant differences in the immature granulocyte frequencies between HD and COVID-19 individuals. However, the proportion of immature granulocytes in moderate and severe COVID-19 donors correlated inversely with the time since onset of symptoms (Fig. S2C). In contrast to previous work (*21*), the total proportion of monocytes (CD14+ HLA-DR+), as well as monocyte subsets (defined by CD14 and CD16), was similar across groups (data not shown). Donors with severe COVID-19 had lower proportions of dendritic cells (DC) compared to moderate disease (p=0.003) and HD (p=0.0374; median percentage in viable CD45+ cells: 0.42% in severe, 0.64% in moderate and 0.49 in HD, Fig. 1A), but not with recovered individuals.

**Fig. 1.**
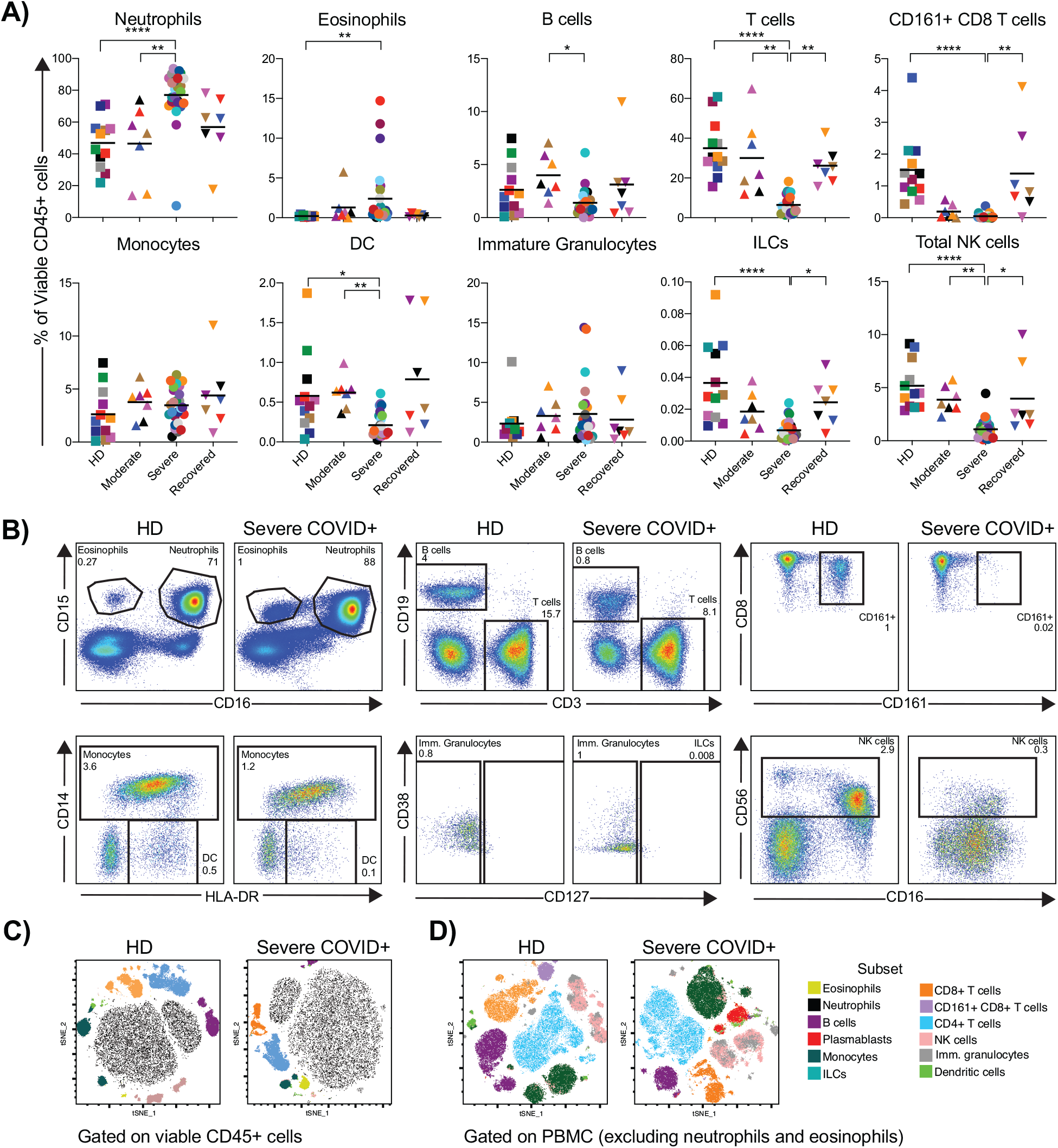
Immune atlas of severe COVID-19. Multiparametric flow cytometry analyses on fresh whole blood after red blood cell lysis characterizing immune cells subsets in healthy donors (HD, n= 12), and moderate (n=7), severe (n=27), and recovered (n=6) COVID-19+ individuals. **A)** Subset frequencies were calculated within the total viable leukocyte CD45+ population. **B)** Dot plots for each immune cell subset in a representative HD and severe COVID-19 individual. Gates within each plot indicate cell subset and corresponding frequency within viable CD45+ cells. Example of parent gates are shown; frequencies were calculated using the specific gating strategies shown in Fig. S1. **C)** Representative examples of the peripheral blood immunologic atlas of a HD and dysregulation within a severe COVID-19 individual. t-distributed stochastic neighbor embedding (t-SNE) analysis of cell subsets gated on total viable CD45+ cells or **D)** PBMC (viable CD45+ cells excluding neutrophils and eosinophils) on a HD and a severe COVID-19 individual. Specific color coding in (**A**) was assigned per individual for cross comparison across Figs. 1-6 and S2-4. Lines on the graphs indicate the median of the group. Differences between groups were calculated using Kruskal-Wallis test with Dunn’s multiple comparison post-test. **** p<0.0001, ***p<0.001, **p<0.01, *p<0.05.

Consistent with previous reports (*7, 8, 22-24*), we observed a relative decrease in the percentages of all lymphocyte subsets (Fig. 1A, B, D). Severe COVID-19 individuals had significantly lower relative proportions of T cells (median frequency within CD45+ cells: 4.5% in severe COVID-19+ and 30.6% in HD; p<0.0001), CD161+ CD8+ T cells (median frequency of CD45+ cells: 0.002% in severe COVID-19 and 1.3% in HD; p<0.0001), innate lymphoid cells (ILCs, median frequency of CD45+ cells: 0.005% in severe COVID-19 and 0.03% in HD; p<0.0001) and natural killer (NK) cells than HD (median frequency of CD45+ cells: 0.95% in severe COVID-19 and 4.5% in HD; p<0.0001). We did not find significant differences in the frequencies of these cell subsets between HDs and moderate or recovered COVID-19 individuals. Within the NK cell lineage, we observed a drastic decrease in the frequencies of both CD56brightCD16- and CD56dimCD16+ NK cells in severe COVID-19 vs. HD (Fig. S2D). In the recovered group, the proportions of T cells, CD161+ CD8+ T cells, ILCs and NK cells were higher than in donors with severe COVID-19 but similar to HDs (median frequencies within viable CD45+ cells: 22% of T cells, 0.1% of CD161+ CD8+ T cells, 0.014% of ILCs, 3.5% of NK cells). The proportions of regulatory CD4+ T cells and circulatory follicular CD4+ T cells were similar across studied groups (Fig. S3A, B). Although we did not observe differences in CD4+ and CD8+ memory T cell subsets between groups (data not shown), we did find a negative correlation with the frequency of central memory T cells (T_CM_) and days since the onset of symptoms (Spearman r= -0.41 p=0.02 for CD4+ T_CM_; Spearman r= -0.61 p=0.0002 for CD8+ T_CM_, Fig. S3C). Given that the neutrophil-to-lymphocyte ratio may be an independent risk factor for severe disease (*25, 26*), we examined the neutrophil:T cell ratio (based on their frequencies within viable CD45+ cells). Individuals with severe COVID-19 had a ratio of 15, while all other studied groups had ratios of less than 2.5. Furthermore, using logistic regression analyses, we did not find any associations between the reported frequencies and comorbidities (pooled together as vascular/metabolic disorders, underlying lung disease and bacterial infections, Table S1). Altogether, these data reveal multiple immunophenotypic abnormalities in severe COVID-19, which are not found in donors with moderate or recovered disease.

### Elevated frequency of plasmablasts, changes in B cell subsets and humoral responses

Although we observed only marginal differences in the proportions of total B cells between the studied groups (Fig. 1), B cell plasmablasts were significantly expanded in severe COVID-19 donors compared to HD (Fig. 1D, Fig. 2A; median frequency within B cells of: 9.7% in severe COVID-19 and 0.48% in HD, p<0.0001). These cells characteristically displayed high levels of Ki-67 and low levels of CXCR5 expression (Fig. S4A). Similar to observations in the immune atlas of recovered COVID-19 donors (*16*), expanded plasmablasts were not found in this group (median frequency with B cells of 0.3% in recovered, p<0.0001 vs. severe donors). The frequency of plasmablasts in individuals with severe COVID-19 did not correlate with age, days since onset of symptoms or the presence of co-morbidities (data not shown), similar to one report based on scRNASeq analyses (*16*).

**Fig. 2.**
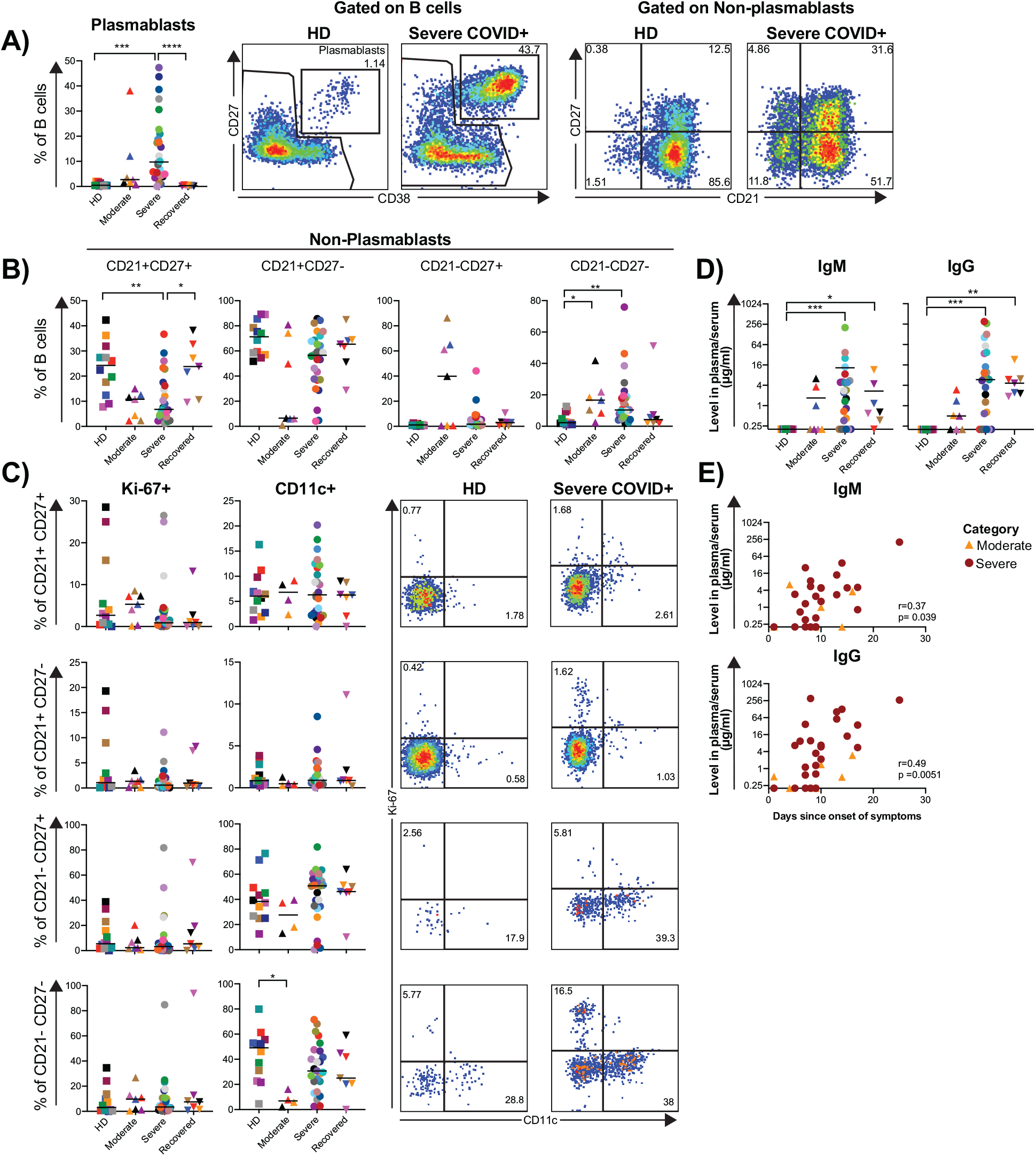
Elevated frequency of plasmablasts, changes in B cell subsets and SARS-CoV-2-specific antibody production in COVID-19 individuals. Multiparametric flow cytometry analyses on fresh whole blood after red blood cell lysis characterizing plasmablast and B cell subset frequencies from HD (n= 12), and moderate (n=7), severe (n=27), and recovered (n=6) COVID-19 individuals. **A), B)** Distribution and representative plots of B cell plasmablasts (defined as CD27+ CD38+ B cells) and non-plasmablast subsets defined by CD21 and CD27 expression in HD (n= 12), and moderate (n=7), severe (n=27), and recovered (n=6) COVID-19 individuals. Numbers inside the plots indicate the subset proportion of the corresponding parent population (within total B cells for plasmablasts, within non-plasmablasts for CD21/CD27 subsets). **C)** Frequencies of CD11c and Ki-67 in non-plasmablast B cell subsets defined in a). Analyses of CD11c are shown for half of the individuals with moderate COVID-19. Plots from a representative HD and severe COVID-19 individual shown. Numbers in each plot indicate the frequency within the parent gate. **D)** Levels of SARS-CoV-2 spike RBD-specific IgM and IgG antibodies in serum or plasma of HD (n= 12), moderate (n=7), severe (n=27), and recovered (n=6) COVID-19 individuals. Antibody measurements were performed by ELISA using plates coated with the receptor binding domain (RBD) from the SARS-CoV-2 spike protein. Sera and plasma samples were heat-inactivated at 56°C for 1 hour prior to testing in ELISA to inactivate virus. Antibody levels were reported as µg/ml amounts relative to the CR3022 monoclonal antibody (recombinant human anti-SARS-CoV-2, specifically binds to spike protein RBD). **E)** Spearman correlations of plasma/serum levels of SARS-CoV-2 RBD-specific IgM (top) and IgG (bottom) and days since onset of symptoms on moderate and severe COVID-19 individuals. Specific color coding was assigned per individual for cross comparison across graphs and Figs. Lines on the graphs indicate the median of the group. Differences between groups were calculated using Kruskal-Wallis test with Dunn’s multiple comparison post-test. **** p<0.0001, ***p<0.001, **p<0.01, *p<0.05.

In the non-plasmablast B cell population, we observed a decrease in the percentage of CD21+CD27+ in moderate and severe groups compared to HD (median frequency of non-plasmablasts of: 24% in HD, 10.8% in moderate disease and 6.7% in severe disease). These proportions were highly significant by nonparametric test of trend (p=0.0008), but only the severe COVID-19 group reached statistical significance vs. HD (p=0.0061, Fig. 2B). Recovered COVID-19 donors had similar levels of CD21+CD27+ non-plasmablasts as the HD group (median of 23.8%). Of note, the frequency of CD21+CD27+ non-plasmablasts was directly correlated with the age of the donors among moderate and severe COVID-19 (Spearman r=0.35, p=0.4, Fig. S4B). In contrast, we observed a significant increase in the proportion of CD21-CD27-non-plasmablasts in moderate (median of 16.6%) and severe (median of 10.4%) COVID-19 individuals compared to HD (median of 2.3%; p=0.0182 and p=0.004, respectively). We next assessed the expression of Ki-67 and CD11c, to determine if any of these subsets were a potential source for the expanded plasmablast population (*27*) (Fig. 2C). We did not observe a larger proportion of cycling Ki-67+ CD21-CD27-B cells in moderate or severe COVID-19 individuals when compared with HD. We also found a reduction in the frequency of CD11c+ cells within CD21-CD27-B cells in donors with moderate COVID-19 compared to HD that was specific to this group (medians of: 6.9% in moderate and 49% in HD; p=0.0162).

Previous work has suggested that the SARS-CoV-2 IgG levels could be associated with disease severity (*12, 28*). With this in mind, and due to the changes observed in B cell subsets, particularly the expansion of plasmablasts in severe COVID-19, we explored the humoral responses in these donors. The levels of total IgG in plasma and serum were equivalent across the groups (Fig. S4C). We then quantified IgM and IgG specific for the spike receptor binding domain (RBD) of the SARS-CoV-2. The levels of both antibodies were significantly higher in the severe and recovered COVID-19 individuals (Fig. 2D). While the frequency of plasmablasts did not correlate with the levels of spike RBD-specific IgM or IgG, there was a positive association between the levels of spike RBD-specific IgM and IgG and time since onset of symptoms (Fig. 2E) in the moderate and severe groups. Together these data indicate an exacerbated plasmablast response in severe COVID-19, as well as the development of a strong SARS-CoV-2-specific humoral response.

### Profound oligoclonal expansion of B cells in severe COVID-19

Having observed the expansion of plasmablasts in severe COVID-19 donors, we sought to determine whether this expansion in severe-COVID-19 resulted from non-specific stimulation. Therefore, we examined the antibody repertoire within samples from randomly selected HD (n=3), moderate COVID-19 (n=3) and severe COVID-19 (n=7) individuals. To sequence antibody heavy chain libraries, we amplified genomic DNA was amplified using primers spanning across nearly the full-length variable (VH) gene sequence and the entire third complementarity determining region (CDR3). After quality control and filtering, the processed antibody heavy chain rearrangements were grouped together into a data set comprising 76 sequencing libraries and 109,590 clones across all 13 individuals (Table S2 and GenBank/SRA PRJNA630455).

To evaluate the clonal landscape, we ranked the proportion of clones within the top ten (1–10), next 90 (11-100), next 900 (100-1,000), and most diverse clones with ranks above 1,000 (1,000+) (Fig. 3A). Donors with severe COVID-19 had an unusually high proportion of large clones comprising the majority of their circulating antibody repertoire, with the fraction occupied by the top 20 ranked clones (D20 measure) the highest compared to the healthy and moderate SARS-CoV-2 infected patients (Fig. 3B, Fig. S5) The D20 rank measure in moderate and severe disease also correlated positively with the plasmablast fraction (Fig. 3C). In many severe COVID-19 individuals we observed very large top copy clones, exceeding the diagnostic thresholds for clinically significant monoclonal B cell lymphocytosis (*29*). These large clones were readily sampled across multiple independently amplified and sequenced libraries (Fig. 3D). Donors M7 and S21 had 91 and 55 clones present in 4 or more sequencing libraries, respectively, in contrast to H4, who had 3 clones in 4 or more libraries (Fig. 3E). Only one HD (H8), an older individual, had large and readily resampled clones, likely reflecting age-dependent narrowing and expansion of the memory B cell repertoire (*30*).

**Fig. 3.**
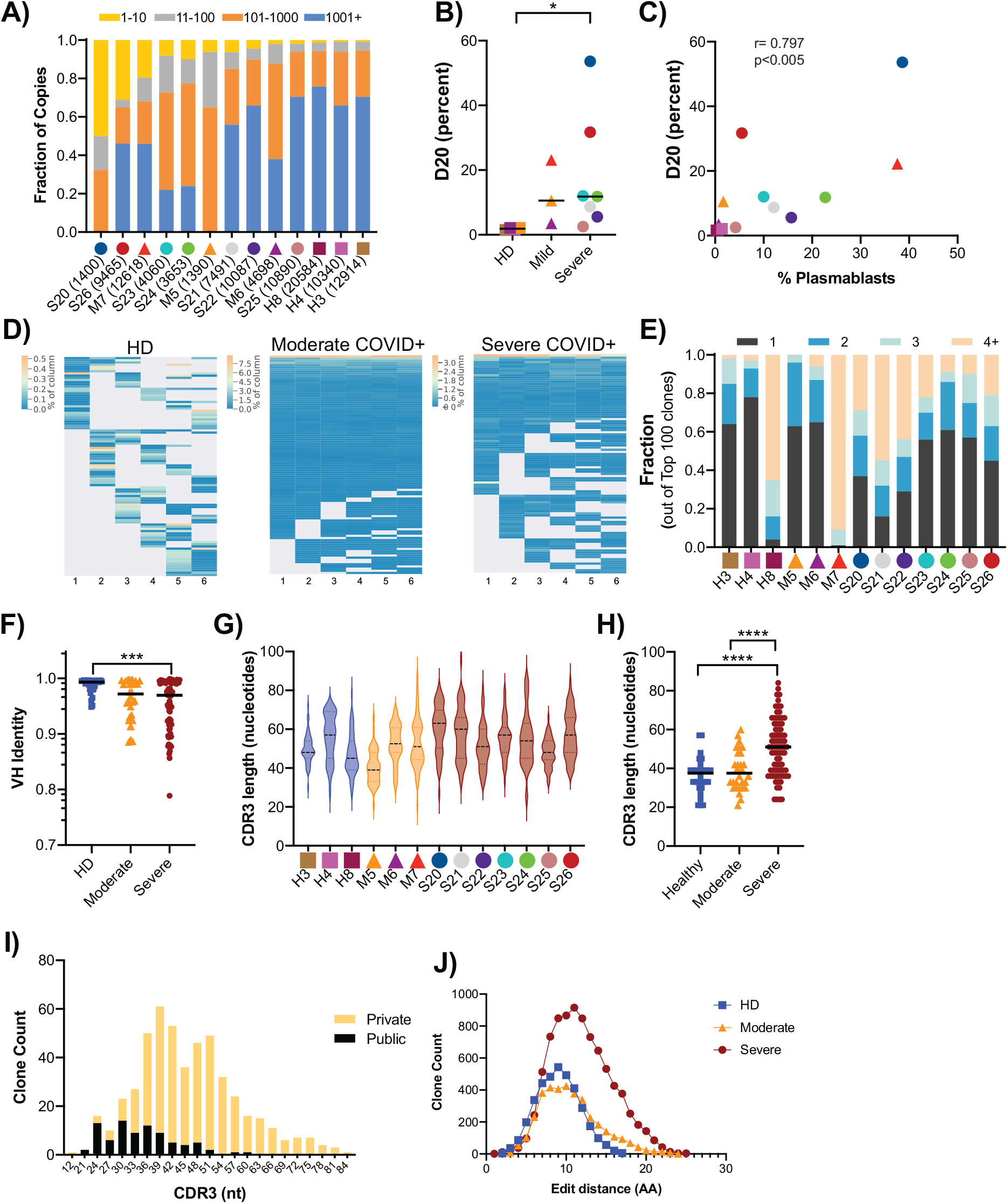
Abundant antibody heavy chain sequences from severe COVID-19 individuals have long, diverse CDR3 sequences and higher levels of somatic hypermutation. **A)** Clone size distribution by sequence copies. For each donor, the fraction of total sequence copies occupied by the top ten clones (yellow), clones 11-100 (grey), 101-1000 (orange) and over 1000 (blue) are shown. Total donor level clone counts are given in parentheses. **B)** Percentage of sequence copies occupied by the top twenty ranked clones (D20) shown for HD (n=3) and COVID-19 patients with moderate (n=3) and severe disease (n=7). **C)** Spearman correlation between the D20 value and the percentage of plasmablasts within the total B cell population. **D)** Examples of the overlap of top 100 copy rearrangements that overlap in at least two sequencing libraries in HD (H4), a moderate COVID-19+ (M7) and a severe COVID-19 individual (S21). Each horizontal string is a rearrangement and each column is an independently amplified sequencing library (see Materials and Methods). Lines are heat mapped by the copy number fraction for a given replicate library. **E)** Clone size estimation based on sampling (presence/absence in sequence libraries). Shown are the fractions of the top 100 clones that are found in 4 or more sequencing libraries, 3 libraries, 2 libraries and 1 library. All donors had six sequencing libraries, except for M5 (four libraries). **F)** Fractional identity to the nearest germline VH gene sequence (1.0 = unmutated) in the top 10 copy number clones of each donor. Each symbol is a clone. **G)** CDR3 length distributions of the top 50 productive rearrangements in each donor. **H)** CDR3 lengths of the top 10 copy number clones (symbols), stratified by condition. **I)** CDR3 length distribution of top 50 clones in COVID-19 donors based on whether they are found in the Adaptive database (public) or not (private). **J)** Distribution of CDR3 amino acid (AA) edit distances of the top 50 copy clones (productive) per donor. Clone pair counts for each edit distance are averaged across all the donors in each disease category. Specific color coding was assigned per individual for cross comparison across graphs and Figs. Lines on the graphs indicate the median of the group. Differences between groups were calculated using Mann-Whitney rank-sum test. **** p<0.0001, ***p<0.001, *p<0.05.

To determine if the antibody heavy chain sequences harbored any evidence of extensive somatic hypermutation (SHM), selective VH gene usage, or defining CDR3 characteristics, we assessed these properties in the top copy clonotypes of each individual. A subset of individuals with severe COVID-19 exhibited higher levels of SHM (Fig. 3F), but other top copy clones in severe COVID-19, moderate COVID-19 and HD were unmutated. To determine if antibodies from COVID-19 individuals exhibited convergent sequence features, we analyzed VH gene usage in all clones of each donor (Fig. S6A). As this analysis did not reveal any consistent increased usage of a specific VH gene in the moderate or severe COVID-19 individuals compared to controls, we reanalyzed the data focusing on the top 200 most frequent clones in each individual (Fig. S6B). Focusing on the most frequently used VH genes, VH genes from different families were used more often in severe COVID-19 donors compared to HD, including VH6-1 (7-fold), VH3-48 and VH3-15 (∼6-fold) and several others (Fig. S6C). We also looked for skewing in VH family usage, which revealed a modest relative increase in the proportion of VH3 family members among COVID-19 individuals compared to HD (Fig. S6D). However, there was considerable inter-individual variation in the usage of VH3 vs. other family members, with some individuals (such as S25) exhibiting substantial skewing towards particular VH families (data not shown).

Given the absence of obvious or uniform VH restriction among COVID-19 individuals, we next analyzed the CDR3 sequences for shared characteristics in the COVID-19 donors. In individuals with severe disease, CDR3 sequences exhibited greater variation in length (Fig. 3G), and were significantly longer among the top copy sequences (Fig. 3H). To determine if the antibody heavy chain sequences from COVID-19 individuals are generated commonly or infrequently, we searched the Adaptive Biotechnologies public database, which consists of 37 million antibody heavy chain sequences (*31*), revealing 3995 matches to the CDR3 amino acid sequences in our dataset. Among the 50 most frequent clones in the COVID-19 individuals, the CDR3 lengths of the matching or “public” clones were shorter than the CDR3 lengths of the non-shared or “private” clones (Fig. 3I), indicating that the top copy clones in COVID-19 with long CDR3 sequences are mostly private. Finally, to determine if there were any collections of clones that harbored similar CDR3 amino acid sequences, we computed the edit distances of all of the amino acid sequences in the top 50 clones of each of the individuals. If there were sequence convergence, we would have expected to find clusters of sequences separated by 3 or fewer amino acids. We found no evidence of co-clustering of CDR3 sequences; rather, over 99% of the edit distances for the severe COVID-19 individuals’ top copy clone pairs were more than 3 amino acids apart (Fig. 3J). Consistent with this finding, alignment of top copy clone CDR3 amino acid sequences from severe COVID-19 individuals revealed highly variable amino acid sequences (Fig. S6E). Taken together, these data show that severe COVID-19 is associated with large, oligoclonal B cell expansions with antibodies enriched for long and divergent CDR3 sequences.

### Innate immune dysregulation in severe COVID-19

Acknowledging the characteristic differences in innate cell subset frequencies in severe COVID-19 individuals (Fig. 1), we further assessed the phenotype of innate immune cells. CD161 has been reported to be a marker of inflammatory monocytes and NK cells (*32–34*). Despite having observed a decreased frequency of CD161+ CD8 T cells (Fig. 1A, D), the frequencies of CD161+ monocytes and CD38+CD161+ NK cells were similar across study groups (Fig. S2E). We next assessed the frequency and expression of CD16 by neutrophils, monocytes, NK cells and immature granulocytes. While the proportions of CD16+ monocytes and immature granulocytes were consistent between groups, severe COVID-19+ individuals had significantly lower circulating CD16+ NK cells in compared with HDs (median percentages of 68% in severe COVID-19 and 85.5% in HD; p=0.0023; Fig. 4A; also observed when analyzing NK cell subsets in Fig. S2D). Furthermore, CD16 expression was significantly lower in neutrophils, NK cells, and immature granulocytes (median fluorescence of CD16 in neutrophils: 7663 in severe and 34458 in HD, p=0.0001; NK cells: 2665 in severe and 10190 in HD; p=0.0017; immature granulocytes: 2728 in severe and 9562 in HD; p=0.0005) in severe COVID-19 (Fig. 4A-F). Downregulation of CD16 in NK cells has been associated with IgG-mediated immune complexes in the context of vaccination (*35*). We did not, however, find significant associations between the frequency or expression of CD16 and IgG levels (Fig. S2F). Although we found a decrease in the frequency of CD16+ monocytes in some severe COVID-19 individuals, this was not consistent amongst the whole cohort (Fig. 1A). The monocyte CD16 expression level tended to decrease with disease severity (median fluorescence intensities of: 5445 in HD, 5235 in moderate and 3619 in severe; p=0.022 by nonparametric test of trend; Fig. 4B). However, monocytes significantly downregulated HLA-DR expression in severe COVID-19 donors compared to moderate disease (p=0.0072) and HD (p=0.021; median fluorescence intensities of 1059 in severe, 4547 in moderate, 5409 in HD; Fig. 4G-H). Similar findings were reported by scRNASeq analysis of severe COVID-19 individuals (*16*) and donors with severe respiratory failure (*36*). In contrast, CD14 expression in monocytes or HLA-DR in other antigen presenting cells (Fig. S2G, H) was consistent across all studied groups. Altogether, these findings indicate a substantial perturbation of the innate immune system in severe COVID-19. Whether this dysregulation is consequence or contributing factor towards COVID-19 severity remains to be defined.

**Fig. 4.**
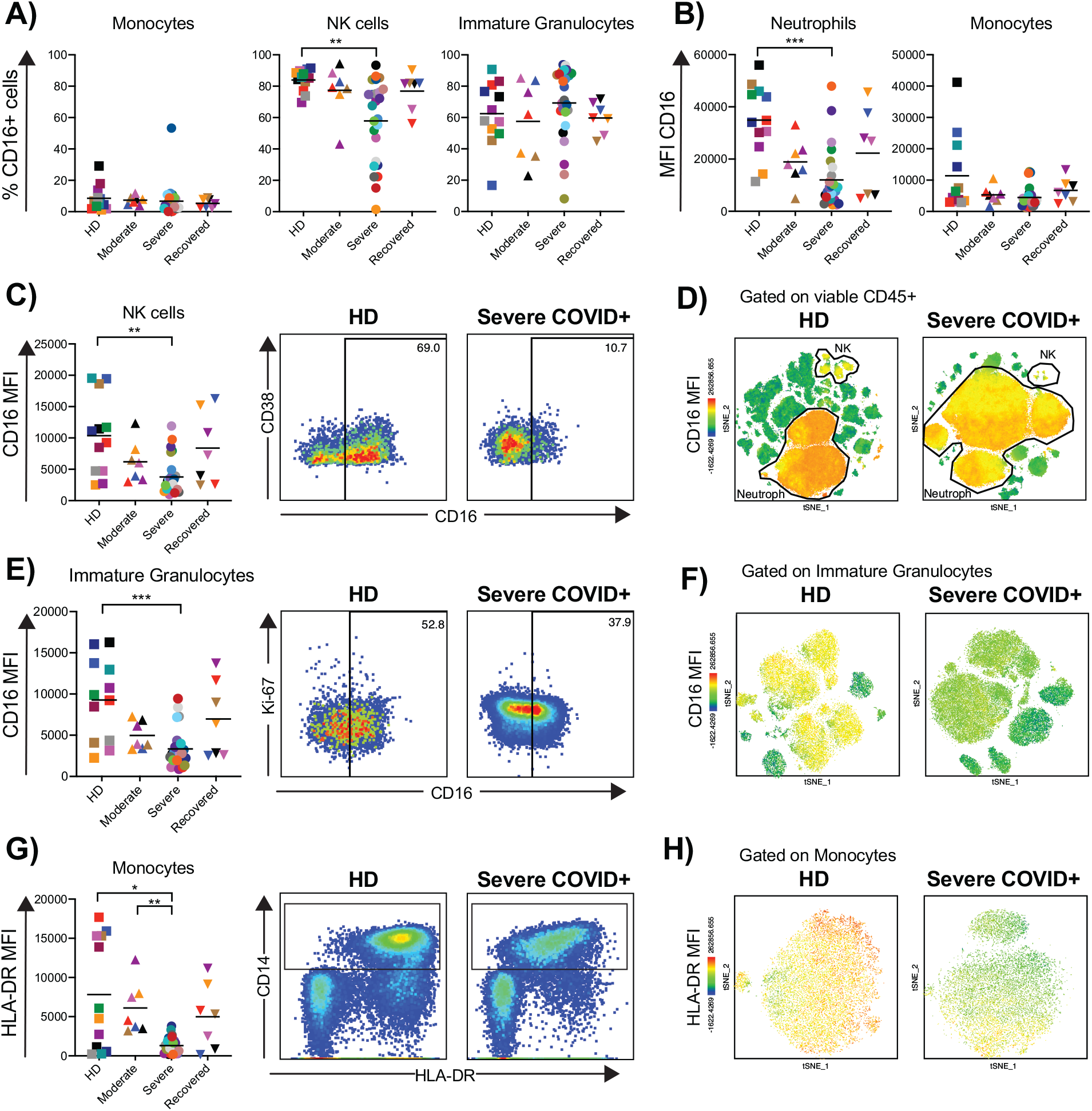
Innate immune dysregulation in severe COVID-19. Multiparametric flow cytometry analyses of fresh whole blood after red blood cell lysis characterizing the expression of CD16 and HLA-DR on innate immune cells from HD (n= 12), moderate (n=7), severe (n=27), and recovered (n=6) COVID-19 individuals. **A)** Proportion of CD16+ cells in monocyte, NK cell and immature granulocyte subsets. **B), C), E)** Median fluorescence intensity (MFI) of CD16 on neutrophil, monocyte, NK cell and immature granulocyte subsets. MFI was calculated within CD16+ cells. Representative dot plots showing CD16 expression in NK cells and immature granulocytes of a HD and a severe COVID-19 individual shown in **C)** and **E)**. The numbers inside the plots indicate the percentage of CD16+ cells in the corresponding parent population. **D), F)** t-SNE analyses of CD16 expression (MFI) in viable CD45+ cells or immature granulocytes, respectively, on a representative HD and a severe COVID-19 individual. **G)** MFI of HLA-DR on monocytes; dot plots of a representative HD and a severe COVID-19 individual shown, with monocyte gate outlined. **H)** t-SNE analyses of monocyte HLA-DR expression (MFI) on a representative HD and a severe COVID-19 individual. Specific color coding was assigned per individual for cross comparison across graphs and Figs. Lines on the graphs indicate the median of the group. Differences between groups were calculated using Kruskal-Wallis test with Dunn’s multiple comparison post-test. ***p<0.001, **p<0.01, *p<0.05.

### Heterogeneous T cell activation in severe COVID-19

T cell activation has been reported in acute respiratory and non-respiratory viral infections (*37–39*). Consistent with recent case reports (*15, 40, 41*), we observed increased activation of both memory CD4+ and CD8+ T cells in severe COVID-19 individuals compared to other study groups (Fig. 5A and B). However, unlike the plasmablast response, heightened T cell activation was not observed in every severe COVID-19 individual and instead demonstrated significant heterogeneity. While overall the frequencies of CD38+ and HLA-DR+ CD38+ memory CD4+ and CD8+ T cells in severe COVID-19 were elevated compared to HD (CD4+, 7.6%, 2.2% vs 2.7%, 0.2%, p=0.009 and p<0.0001, respectively; CD8+, 9.2%, 3.9% vs. 0.6%, 0.09%; p<0.0001 for both cases), we did not find statistically higher Ki-67+ CD4+ or CD8+ T cells in COVID-19 individuals compared to HD. However, a subset of severe COVID-19 donors clearly had increased levels of Ki-67+ CD4+ and CD8+ T cells, reaching as high as ∼25% in some individuals. The frequency of PD-1+ memory CD4+ T cells (44.3% in severe and 25.7% in HD, respectively; p=0.0084), but not CD8+ T cells, was also higher in the severe COVID-19 group compared to the HD group. For all measures, CD4+ and CD8+ T cell activation in recovered donors was equivalent to the HD group. Of note, the proportion of PD-1+ memory CD4+ T cells, but not of PD-1+ CD8+ T cells, in moderate or severe COVID-19 correlated with donor age (Fig. S3D). In addition, the frequencies of HLA-DR+ CD38+ CD4+ and CD8+ T cells correlated with the proportion of plasmablasts in moderate and severe COVID-19 individuals (r=0.5011 p=0.0022, and r=0.4722 p=0.0042, respectively, Fig. 5C).

**Fig. 5.**
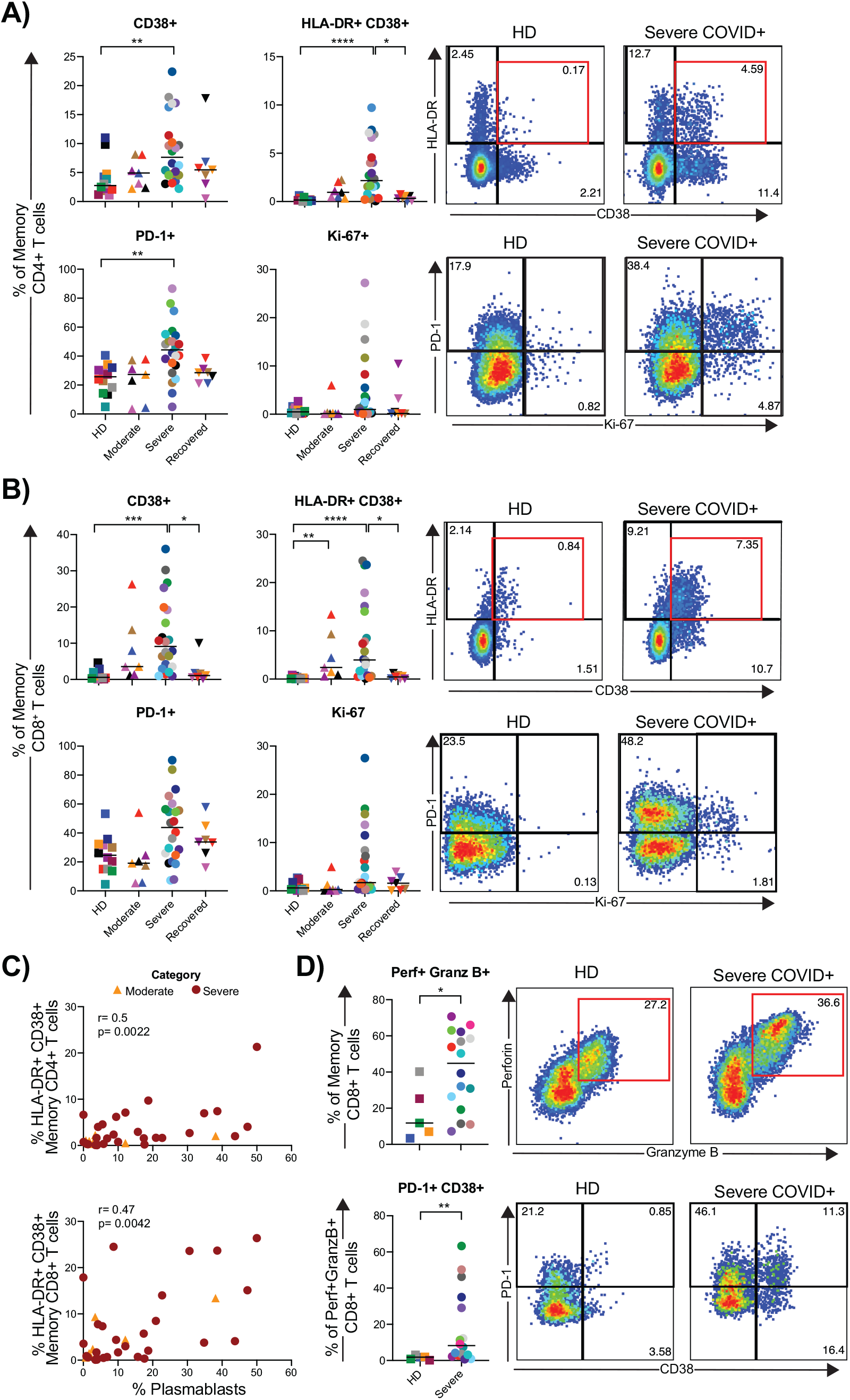
Heterogeneous T cell activation in severe COVID-19. Multiparametric flow cytometry analyses on fresh whole blood after red blood cell lysis characterizing immune cells subsets in HD (n= 12), moderate (n=7), severe (n=27), and recovered (n=6) COVID-19 individuals was performed to assess the percentage of activated memory T cells. Frequencies of CD38+, HLA-DR+CD38+, PD-1+ and Ki67+ in **A)** CD4+, and **B)** CD8+ memory T cells (excluding naïve CCR7+ CD45RA+, detailed gating strategy shown in Fig. S1). **C)** Spearman correlations between the frequencies of HLA-DR+ CD38+ CD4+ or CD8+ memory T cells and plasmablasts in donors with moderate (orange triangles) or severe COVID-19 (dark red circles). **D)** Frequency of cytotoxic memory CD8+ T cells. Multiparametric flow cytometry analyses were performed on freshly isolated PBMC from HD (n=5) and severe (n=16) COVID-19 individuals to quantify the frequency and phenotype of cytotoxic (as defined by perforin and granzyme B expression) CD8+ T cells, and proportion of cytotoxic CD8+ T cells expressing PD-1 and CD38. Plots for a representative HD and a severe COVID-19 individual are shown. Numbers inside the plots indicate the frequency within the corresponding parent population. Specific color coding was assigned per individual for cross comparison across graphs and Figs. Lines on the graphs indicate the median of the group. Differences between groups were calculated using Kruskal-Wallis test with Dunn’s multiple comparison post-test and Mann-Whitney rank-sum test. **** p<0.0001, ***p<0.001, **p<0.01, *p<0.05.

We further quantified the proportion of cytotoxic CD8+ T cells (defined as perforin+ granzyme B+ memory CD8+ T cells, Fig. 5D) in a subset of HD and severe COVID-19 individuals. Due to limited samples, we did not include the moderate or recovered COVID-19 groups for this analysis. We found a significantly higher proportion of cytotoxic CD8+ T cells in severe COVID-19 than in HD (median frequency within memory CD8+ T cells of 48.7% and 27.2%, respectively; p=0.048). The frequencies of T-bet+ cells, as well as the levels of expression (measured by median fluorescence intensity) of perforin+ and granzyme B+ cells within the cytotoxic memory CD8+ T cell subset were similar between groups (Fig. S3E-F). Cytotoxic CD8+ T cells from severe COVID-19 donors also had an increased proportion of cells expressing CD38 or co-expressing PD-1 and CD38 compared to HD (medians of 8.2% and 1.8%, respectively; p=0.0082; Fig. 5D and Fig. S3G). These data indicate a heightened status of immune activation and frequency of cytotoxic CD8+ T cells during severe COVID-19, not observed in moderate or recovered disease.

### Distinctive severe COVID-19 immunophenotype

Finally, we performed an unbiased analysis to determine if the immune cells in severe COVID-19 disease cohort could be differentiated from the healthy, moderate, and recovered cohorts. We included all analyzed immune phenotype parameters described thus far, including the expression of activation markers within specific CD4+ and CD8+ T cell memory subsets (data not shown). We scaled all flow cytometry generated data using z-score, and performed hierarchical clustering (Fig. 6A). From this analysis, the data from 21/28 of the severe COVID-19 patients co-localized to a distinct cluster within the hierarchical tree. We further analyzed these data by principal component analysis, where we again found selective clustering of individuals with severe COVID-19 (Fig. 6B). The top parameters driving the clustering of the severe COVID-19 were associated with T cell activation in CD4+ and CD8+ T cell memory subsets, frequency of plasmablasts and frequency of neutrophils (Table S3), also evidenced in the heat map shown in Fig. 6A. Independent analyses of the severe COVID-19 group did not produce separate clustering, likely due to reduced sample number. However, it is clear from the heatmap analysis that distinct patterns within the severe COVID-19 disease cohort may be present that further subdivide these individuals into different subgroups. Taken as a whole, our analysis reveals a characteristic immune phenotype in severe COVID-19, distinct not only from HD but also from other COVID-19 individuals with moderate or recovered disease.

**Fig. 6.**
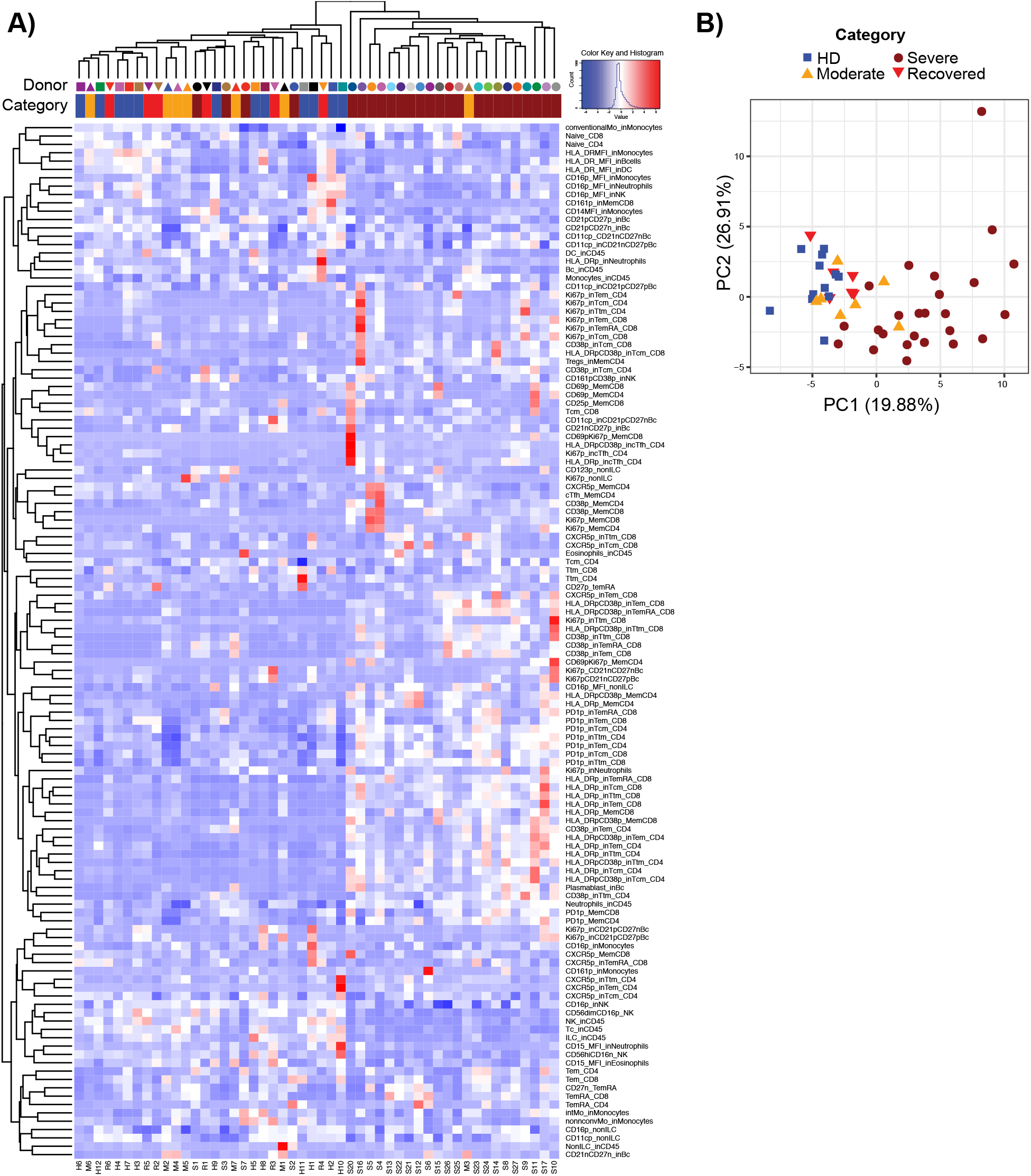
Unbiased analyses of immunophenotyping reveals selective clustering of severe COVID-19 individuals. **A)** Heatmap of flow cytometric analyses of HD (n= 12), moderate (n=7), severe (n=27), and recovered (n=6) COVID-19 patients. Data are shown in z-score scaled values. Shape and color coding correspond to data shown in Figs. 1-6. H, HD; M, moderate COVID-19; S, severe COVID-19; R, recovered COVID-19. **B)** Principal component analysis generated using all flow cytometric data from **A)**.

## Discussion

Devising therapeutic strategies to treat SARS-CoV-2 infection remains challenging, due to both the complexity of the clinical manifestations and an overall lack of understanding of severe COVID-19 immunopathogenesis. Reports on single individuals, studies with small patient numbers of varying disease stages, or focused analyses on limited immune phenotypes have generated valuable information, but have fallen short of providing a comprehensive immunophenotypic atlas of severe COVID-19. Here, we sought to define immune perturbations of COVID-19 in moderate and severe disease using an unbiased approach designed to simultaneously capture changes in the predominant granulocyte and lymphocyte populations. We found profound changes in multiple leukocyte populations selectively in severe disease that provides both novel and confirmatory insights into the immunopathogenesis of severe COVID-19, including pronounced effects on neutrophils, monocytes, NK cells, and B and T lymphocytes.

Modulation of innate immune cells manifested in a number of ways, including broad downregulation of CD15 and CD16 on neutrophils, as well as CD16 downregulation on NK cells, immature granulocytes and monocytes. Retrospective clinical metadata studies have identified an elevated neutrophil:lymphocyte ratio in severe COVID-19, a finding we confirm here (*25*). It is unclear whether CD15 and CD16 downregulation marks an activated or refractory state. On NK cells, CD16 downregulation has been associated with NK cell maturation and development (*42*), as well as with activation and target cell engagement, resulting in antibody derived cell cytotoxicity and TNF-alpha secretion. Alternatively, downregulation of CD16 after interaction with IgG-immune complexes also may prevent excessive immune responses after influenza vaccination (*35, 43*). Although it did not reach statistical significance between groups, we also observed lower CD16 expression on monocytes in severe COVID-19 individuals compared to HD. Similar to its suggested role in NK cells, regulation of CD16 in monocytes promotes TNF-alpha production upon cross-linking by immune complexes and phagocytosis through IgG (*44*). Analogous changes in phenotype of innate immune cells have been reported in other conditions and infectious diseases (*45–47*). The implications of the observed changes in the expression of CD15 in neutrophils, as well as CD16 across subsets during severe COVID-19 and their potential role as indicators of redistribution to the lungs, link with function and response, as well as diagnostic and prognostic significance (*48–50*), requires additional exploration.

One of our most striking findings was a profound expansion of plasmablasts during severe COVID-19, in some patients rivaling or exceeding that observed in acute hantavirus, dengue and Ebola infections or chronic inflammatory conditions such as systemic lupus erythematosus (*38, 51–54*). One recent study suggested that COVID-19+ individuals in critical condition show extrafollicular B cell activation (*55*). The increase in the plasmablast frequency we observed directly correlated with an oligoclonal expansion of antibody clones within the overall B cell repertoire, suggesting that many of these large clonal expansions reside within the plasmablast pool. Remarkably, in some severe COVID-19 individuals a single clone could account numerically for the entire plasmablast population. Only one individual with moderate disease displayed this marked plasmablast expansion, the majority harboring smaller clones with more diverse repertoires. The antibody sequences of the largest B cell clones in the severe COVID-19 individuals were surprisingly variable in terms of SHM levels, but consistently had long CDR3 regions compared to donors with moderate COVID-19 and HD. B cells harboring antibodies with long CDR3 sequences are often multi-reactive and counter-selected during B cell development (*56*), which may suggest a contribution of longer CDR3 sequences as part of severe COVID-19 immunopathology.

In line with a recent report (*57*), we did not observe clear sequence convergence of VH genes amongst all the severe COVID-19 individuals, but VH3 family members were enriched in some individuals. CDR3 sequences from individuals with severe COVID-19 had higher edit distances than individuals with mild disease or HD. While their size, somatic mutation status and association with the plasmablast fraction are suggestive of active participation in the immune response to SARS-CoV-2, it is unknown if these clones can recognize the virus, confer protection, or contribute to immunopathology. Future comparisons of our data to antibodies of known specificity may provide important insights into the dynamics of antibody responses in different phases of the illness and may reveal important differences between antibodies produced in the context of moderate vs. severe disease.

T cell activation is typically observed during acute viral infections (*58–60*), and as expected (*15, 18*) we observed increased activation of both CD4+ and CD8+ T cells in severe COVID-19 that correlated with the plasmablast frequency. However, T cell activation was very heterogeneous across the severe COVID-19 patients, being equivalent to baseline in some while reaching up to ∼25% of memory CD8+ T cells in others. This heterogeneity is relatively unusual compared to the symptomatic phase in other acute infections in humans, such as HIV, EBV, HCMV, HBV, and Ebola, where activation is uniformly detectable but to varying, and sometimes much higher, degrees (*61–64*). However, given the degree of lymphopenia observed in the severe COVID-19 patients, it is possible that activated T cells are migrating to, or sequestered in, the lung in response to the virus (*23, 65–68*), making it unclear if T cell activation is found in other sites as suggested by case study reports (*6, 69*). We also observed a marked reduction in the frequency of CD161+CD8+ T cells in donors with severe COVID-19. This subset is composed primarily of mucosal-associated invariant T cells (MAIT) cells (< 95%) (*70*) and a small subset of IL-17 secreting cells (Tc17) (*71*). During viral infections, both MAIT and Tc17 cells can become activated and migrate to infection sites (*71, 72*). Critically ill COVID-19 individuals were recently shown to have a profound decrease in circulating MAIT cells paralleled with their presence in airways (*73*). As such, the reduction of CD161+ CD8+ T cells in periphery found here could be indicative of cell sequestration to the lungs, potentially exacerbating tissue inflammation.

Many of the immunological characteristics of severe COVID-19 share features of sepsis-associated immune dysregulation, yet others are more specific for an acute viral infection. Decreased expression of CD16 on neutrophils, monocytes, and immature granulocytes and decreased expression of HLA-DR in monocytes has been associated with sepsis and sepsis outcome (*36, 74–78*). However, expansion of plasmablasts and activated T cells is common to typical acute viral infections, not sepsis. Severe COVID-19 is a distinct clinical and immune sepsis subphenotype, and the immune dysregulation may necessitate targeted strategies to effectively manage clinical care. To this end, the immunological analysis strategy that we presented readily differentiated those with severe COVID-19 compared to HD, moderate cases, and recovered cases. Longitudinal studies to determine whether early detection of the immunological perturbations that we have defined here predicts severe disease trajectory, even when patients exhibit only asymptomatic or moderate disease could provide crucial insight into the development of effective therapeutic interventions to ameliorate severe COVID-19.

## Acknowledgements

The authors would like to thank all blood donors, their families and surrogates, as well as the medical personnel in charge of patient care. This work was supported by the University of Pennsylvania Institute for Immunology Glick COVID-19 research award (MRB); NIH HL137006 and HL137915 (NJM); Mentored Clinical Scientist Career Development Award from the National Institute of Allergy and Infectious Diseases K08 AI136660 (LV); NIH UM1-AI144288 and P30-CA016520 (WM, AMR, ETLP); NIH AI105343, AI115712, AI117950, AI108545, AI082630 and CA210944 (EJW). NJM reports funding to her institution from Athersys, Inc, Biomarck Inc, and the Marcus Foundation for Research. EJW is supported by the Parker Institute for Cancer Immunotherapy which supports the Cancer Immunology program at the University of Pennsylvania. We thank Florian Krammer (Mt. Sinai) for providing the SARS-CoV-2 spike RBD expression plasmid used to produce antigen for IgM/IgG ELISAs.

## Author Contributions

LK-C, MBP conceptualized, designed, conducted and analyzed all flow cytometry and total IgG quantification experiments. WM conducted IgH sequencing experiments. WM, AMR and ETLP analyzed sequencing data. LK-C, MBP, DM, AEB, ARG, AP, JK, and NH processed blood samples. NJM, NSM, TKJ, ARW, CAGI, RA, OK, LV, SA, LB and JD conducted donor recruitment and collected all relevant clinical information. SG, MEW, CPA, MJB, ECG, EMA and EZM performed IgG and IgM quantification, supervised by SEH. LK-C, MBP, ETLP and MRB wrote the paper. MRB, NJM and EJW supervised the study.

## Data and materials availability

All data associated with this study are present in the paper or the Supplementary Materials. The immunoglobulin heavy chain sequencing data is being submitted in an AIRR-compliant manner to SRA under PRJNA630455.

## Supplementary Materials

Materials and Methods

Fig. S1 Gating strategy used for flow cytometric analyses of immune cell subsets.

Fig. S2. Extended innate immune subset characterization and phenotype during COVID-19 infection.

Fig. S3. Extended T cell phenotype and activation during COVID-19 infection.

Fig. S4. Extended B cell phenotype and total IgG measurements in COVID-19.

Fig. S5. Abundance of the top 20 clones in each donor.

Fig. S6. Heavy chain variable (VH) gene and CDR3 usage. Fig. S4. Extended immune subset characterization and phenotype during COVID-19 infection.

Tables S1. Detailed clinical characteristics of individuals with moderate and severe COVID-19.

Table S2. Antibody heavy chain gene rearrangement metadata.

Table S3. Rotation table extracted from PCA.

### Materials and Methods

#### Blood donors

Inpatient donor admitted to the Hospital of the University of Pennsylvania with a SARS-CoV-19 positive result were screened and approached for informed consent within three days of hospitalization. Recovered donors with a prior positive SARS-CoV-19 test and healthy donors were recruited initially by word of mouth, and subsequently through a centralized University of Pennsylvania resource website for COVID-19-related studies. All participants or their surrogates provided informed consent in accordance with protocols approved by the regional ethical research boards and the Declaration of Helsinki. Peripheral blood was collected from all donors. For inpatients, clinical data were abstracted from the electronic medical record into standardized case report forms. ARDS was categorized in accordance with the Berlin definition reflecting each subject’s worst oxygenation level and with physicians adjudicating chest radiographs (*1*). APACHE III scoring was based on data collected in the first 24 hours of ICU admission or the first 24 hours of hospital admission for subjects who remained on an inpatient unit. Clinical laboratory data was collected from the date closest to the date of research blood collection.

#### Sample Processing

Peripheral blood samples processed within 3 hours of collection. After plasma separation, 1 ml of whole blood was separated for staining and the remaining volume was used for PBMC isolation using SepMate tubes (StemCell Technologies, Vancouver, Canada) following manufacturer’s instructions.

#### Whole blood and PBMC staining

Flow cytometry experiments were performed on whole blood or freshly isolated PBMC. For whole blood stains, leukocytes were obtained after lysis of red blood cells using ACK buffer (Thermofisher, Waltham, MA) during 5 minutes followed by a wash with R10 media (RPMI-1640 supplemented with 10% FBS, 2 mM L-glutamine,100 U/ml penicillin, and 100 mg/ml streptomycin). After washing with phosphate-buffered saline (PBS), cells (whole blood derived leukocytes or PBMC) were prestained for the chemokine receptor CCR7 for 10 min at 37°C 5% CO_2._ All following incubations were performed at room temperature. Cells were stained for viability exclusion using Live/Dead Fixable Aqua for 10 minutes, followed by a 20-minute incubation with a panel of directly conjugated monoclonal antibodies and Trustain diluted in equal parts of fluorescence-activated cell sorting (FACS) buffer (PBS containing 0.1% sodium azide and 1% bovine serum albumin) and Brilliant stain buffer (BD Biosciences, San Jose, CA). The cells were washed in FACS buffer and fixed/permeabilized using the FoxP3 Transcription Factor Buffer Kit (eBioscience, San Diego, CA), following manufacturer’s instructions. Intracellular staining was performed by adding the antibody cocktail prepared in 1X permwash buffer for 1 hour at 37°C. Stained cells were washed and fixed in PBS containing 4% paraformaldehyde (Sigma-Aldrich, St. Luis, MO), and stored at 4°C in the dark until acquisition.

All flow cytometry data were collected on a BD FACSymphony A5 cytometer (BD Biosciences). Data were analyzed using FlowJo software (version 10.6.2, Tree Star, Ashland, OR).

#### Antibodies

The following antibodies were used: CD69 PE-Cy5 (clone FN50), PD-1 BV421 (clone EH12.2H7), CCR7 APC-Cy7 (G043H7), CD19 BV785 (clone HIB19), CD27 BV650 (clone 0323), CD56 BV570 (clone HCD56), CD16 BV711 (clone 3G8), CD21 PE-Cy7 (clone BU32) and Perforin APC (clone B-D48) from Biolegend (San Diego, CA); CD11c PE-Cy5.5 (clone N418) from eBioscience; CXCR5 BV750 (clone RF8B2), CD3 BUV805 (UCHT1), CD45 AF700 (clone HI30), CD127 PE-CF594 (clone HIL-7R-M21), CD25 BUV737 (clone 2A3), CD8 BUV496 (clone RPA-T8), HLA-DR BV605 (clone G46-6), CD123 PE (clone 9F5), CD38 BUV661 (clone HIT2), CD14 BV480 (clone MP9), CD45RA BUV563 (HI100), CD4 BB790 (clone SK3), CD15 FITC (clone HI98), CD103 BB700 (clone Ber-ACT8), CD161 APC (clone DX12), Ki-67 BUV395 (clone B56) and Granzyme B FITC (clone GB11) from BD Biosciences (San Diego, CA). The Live/Dead Fixable Aqua Dead Cell Stain Kit (Invitrogen) was used for viability exclusion, and Human Trustain FcX (Biolegend) was used to prevent unspecific binding.

#### Quantification of total plasma/serum IgG by Cytometric Bead Array (CBA)

Total IgG was measured using a Hu Total IgG CBA Flex Set Bead (BD Biosciences) on plasma or serum samples following manufacturer’s protocol.

#### Enzyme-linked immunosorbent assay (ELISA) for SARS-CoV-2-specific antibody quantification

ELISAs were completed to measure antibodies against the SARS-CoV-2 receptor binding domain (RBD) protein as previously described (*2*). Plasmids encoding the SARS-CoV-2 RBD were provided by Florian Krammer (Mt. Sinai) (*3, 4*). SARS-CoV-2 RBD proteins were produced in-house in 293F cells and purified using Ni-NTA resin (Qiagen, Germantown, MD). ELISA plates (Immulon 4 HBX, Thermo Scientific) were coated with 50 μL per well of recombinant protein diluted in PBS to a final concentration of 2μg/mL and plates were incubated overnight at 4°C. The next day, ELISA plates were washed 3 times with PBS containing 0.1% Tween-20 (PBS-T) and were blocked with PBS-T supplemented with 3% non-fat milk powder for 1 hour at room temperature. Sera and plasma samples were first heat-inactivated at 56°C for 1 hour and then serially diluted in 2-fold in PBS-T supplemented with 1% non-fat milk powder (dilution buffer) starting at a 1:50 dilution. ELISA plates were washed 3 times with PBS-T before the addition of 50 μL of diluted serum and were incubated for 2 hours at room temperature. Goat anti-human IgG-HRP (Jackson ImmunoResearch Laboratories, West Grove, PA) was diluted 1:5000 and goat anti-human IgM-HRP (SouthernBiotech, Birmingham, AL) was diluted 1:1,000 in dilution buffer. After ELISA plates were washed 3 times with PBS-T, 50 μL of secondary antibodies were added to each well and plates were incubated for 1 hour at room temperature. ELISA plates were washed 3 times with PBS-T and were developed for 5 mins at room temperature with 50 μL per well of SureBlue TMB substrate (KPL). The reaction was stopped by acidification with the addition of 25 μL of 250 mM hydrochloric acid and optical density (OD) readings at 450 nm were obtained using the SpectraMax 190 microplate reader (Molecular Devices, San Jose, CA). An anti-SARS-CoV S therapeutic monoclonal antibody (CR3022) was included on each plate and serum/plasma antibody levels were reported as relative μg/mL amounts. Plasmids to express the CR3022 monoclonal antibody were provided by Ian Wilson (Scripps).

#### Antibody heavy chain sequencing

DNA was extracted from blood using Gentra Puregene Blood Kit (Qiagen). Immunoglobulin heavy-chain family-specific PCRs were performed on genomic DNA samples using primers in FR1 and JH as described previously (*5*). Six biological replicates at 400 ng input DNA per replicate were run on all subjects except for subject M5 (4 replicates and 63.5 ng DNA/replicate) and S20 (6 replicates at 333.7 ng DNA/replicate). Sequencing was performed in the Human Immunology Core Facility at the University of Pennsylvania. Illumina 2 × 300-bp paired-end kits were used for all experiments (Illumina MiSeq Reagent Kit v3, 600-cycle, Illumina MS-102-3003).

#### Antibody heavy chain sequence analysis

##### Quality Control, gene identification and clonal inference

Sequencing data were quality controlled with pRESTO (*6*), using a similar protocol described previously (*7*). DNA was chosen for this analysis because it provided a parsimonious means of evaluating the B cell repertoire, with one template per cell, and because replicate sequencing libraries could be used to provide rigorous clone size estimates (*7*). Briefly, paired reads were assembled using default parameters, sequences that had an average quality score less than 30 were excluded, ends of each read which had an average quality score less than 30 within a window of 20 bases were trimmed, sequences shorter than 100 nucleotides were excluded, and bases with a quality score less than 30 were masked with an *N*. Sequences with ten or more *N*s were then discarded. Sequences were annotated with IgBLAST, (*8*) and imported into ImmuneDB v0.29.9 (*9*) for further processing and data visualization. To group related sequences together into clones, ImmuneDB hierarchically clusters sequences with the same VH gene, same JH gene, same CDR3 length, and 85% identity at the amino acid level within the CDR3 sequence (*5*). Clones with consensus CDR3 sequences within 2 nucleotides (*10*) of each other were further collapsed to account for incorrect gene calls.

##### Data Visualization

Data were exported from ImmuneDB for downstream analysis. pandas v1.0.0 was used for data manipulation, seaborn v0.10.0 and Prism v8.4.0 were used for graphing, scipy v1.4.1 for statistical testing, and python-Levenshtein v0.12.0 was used for edit distance calculations. Edit distance was calculated using an unweighted Levenshtein distance (*11*). The edit distance between two CDR3 strings is the number of insertions, deletions, or substitutions required to convert one string into the other. The somatic hypermutation (SHM) of a given clone was determined by comparing every unique sequence in the clone to the most similar VH germline gene sequence. SHM is defined as the percentage of mismatching nucleotides compared to the closest corresponding germline gene. Only the VH portion, not the CDR3 or J-region, was included in the SHM calculation. CDR3 sequence analysis was performed using Geneious Prime 2020.1.2.

#### Statistical Analysis

All statistical analyses were performed using GraphPad Prism (version 8.4.2 GraphPad Software, La Jolla California USA) and R software (URL http://www.R-project.org/). Kruskal-Wallis ANOVA with Dunn’s multiple comparison tests or Mann-Whitney tests were used to compare between groups, or one-way ANOVA with non-parametric test for trend, as appropriately indicated. Non-parametric Spearman correlations or simple logistic regression analyses were used to determine associations between analyzed parameters.

---

**Fig. S1.**
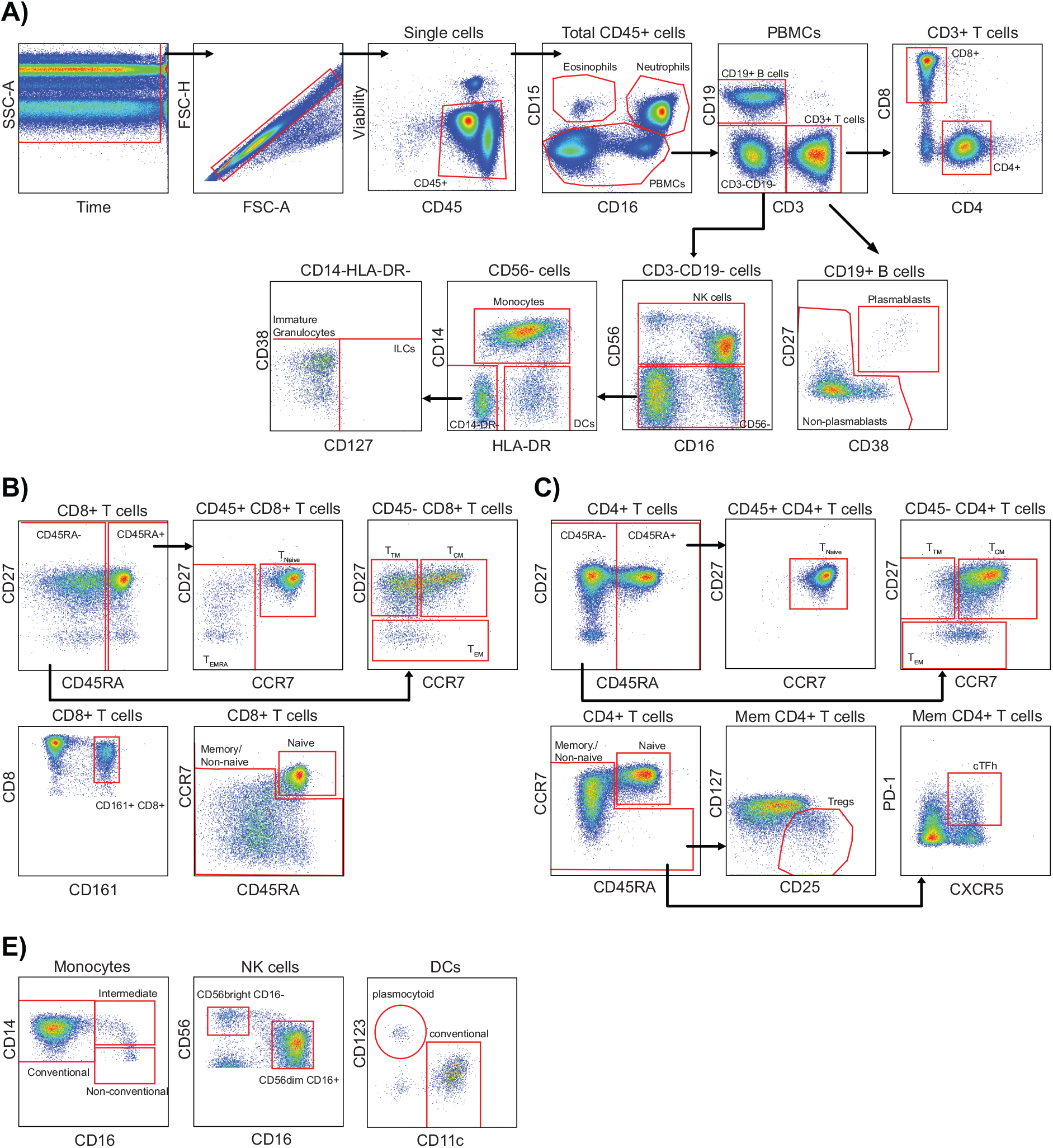
Gating strategy used for flow cytometric analyses of immune cell subsets. Representative example of a HD is shown. **A)** Identification of eosinophils, neutrophils, B cells (plasmablasts and non-plasmablasts), T cells, NK cells, monocytes, dendritic cells (DCs), innate lymphoid cells (ILCs) and immature granulocytes in whole blood. Cleaning gates were performed for each subset before calculating frequency within viable CD45+ cells, and phenotype characterization (neutrophils were further cleaned for the expression of CD4, CD8, CD14, CD19; T cells were cleaned for CD14 and CD15; B cells were cleaned for CD3, CD14, CD15 and CD56; CD3-CD19-cells were cleaned for CD3 and CD15). **B)** Characterization of CD8+ T cell subsets as defined by expression of CD27, CD45RA and CCR7. CD161+ CD8+ T cells were analyzed within the whole CD8+ T cell population. Expression of activation markers was also determined in the whole memory/non-naïve CD8+ T cell subset. **C)** Characterization of CD4+ T cell subsets as defined by expression of CD27, CD45RA and CCR7. Expression of activation markers and other subsets were also determined within the whole memory/non-naïve CD4+ T cell subset. Regulatory CD4+ T cells were defined as CD127low CD25+, and circulating follicular CD4+ T cells as CXCR5+ PD-1+ within the memory subset. **D)** Identification of monocyte, NK and DC subsets. T_CM_, central memory T cells; T_EM_, effector memory T cells; T_TM_, transitional memory T cells; T_EMRA_, CD45RA+ effector memory T cells; Tregs, regulatory CD4+ T cells; cTfh, circulating follicular CD4+ T cells.

**Fig. S2.**
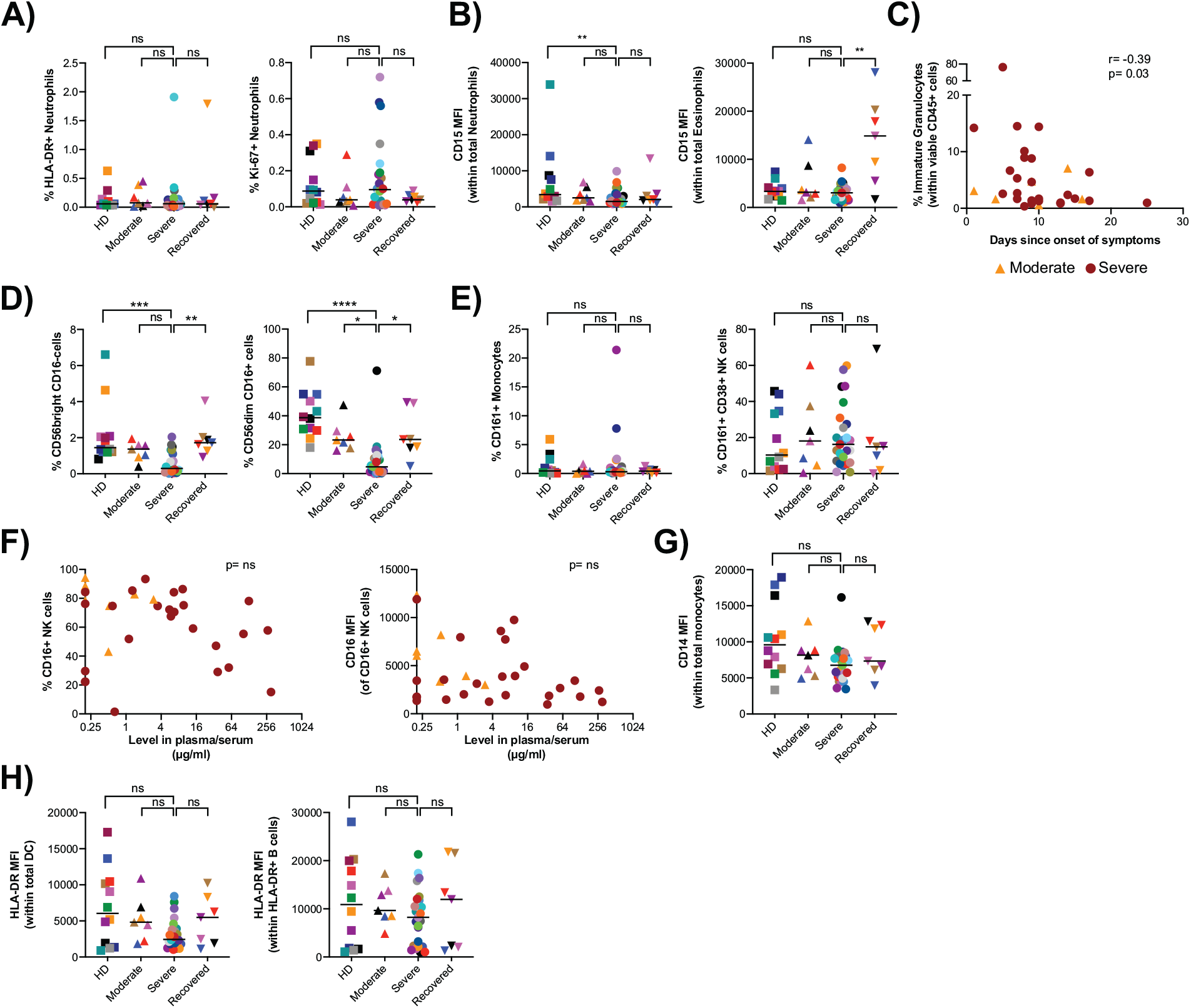
Extended innate immune subset characterization and phenotype during COVID-19 infection. Multiparametric flow cytometry analyses on fresh whole blood after red blood cell lysis characterizing immune cells subsets in HD (n= 12), moderate (n=7), severe (n=27), and recovered (n=6) COVID-19+ individuals was performed. **A)** Frequency of activated or cycling neutrophils, measured by the frequency of HLA-DR+ or Ki-67+ cells. **B)** Mean fluorescence intensity (MFI) of CD15 in neutrophils and eosinophils. **C)** Spearman correlation of the frequency of ILCs and days since onset of symptoms in moderate (orange triangles) and severe COVID-19+ individuals (dark red circles). **D)** Percentages of CD56bright and CD56dim NK cell subsets. Frequencies within parent population are shown (CD3-CD19-cells). **E)**Proportion of inflammatory monocytes or NK cells (gated in total CD56+ NK cells), defined by the single expression of CD161+ or co-expression of CD161 and CD38, respectively. **F)** Spearman correlation of the percentage of CD16+ and expression (MFI of CD16+ cells) and plasma/serum RBD-specific IgG levels in moderate (orange triangles) and severe COVID-19+ individuals (dark red circles). **G)** MFI of CD14 in monocytes. **H)** MFI of HLA-DR+ dendritic cells and B cells (non-plasmablasts). Specific color coding was assigned per individual for cross comparison across graphs and Figs. Lines on the graphs indicate the median of the group. Differences between groups were calculated using Kruskal-Wallis test with Dunn’s multiple comparison post-test. **** p<0.0001, ***p<0.001, **p<0.01, *p<0.05, ns, not significant.

**Fig. S3.**
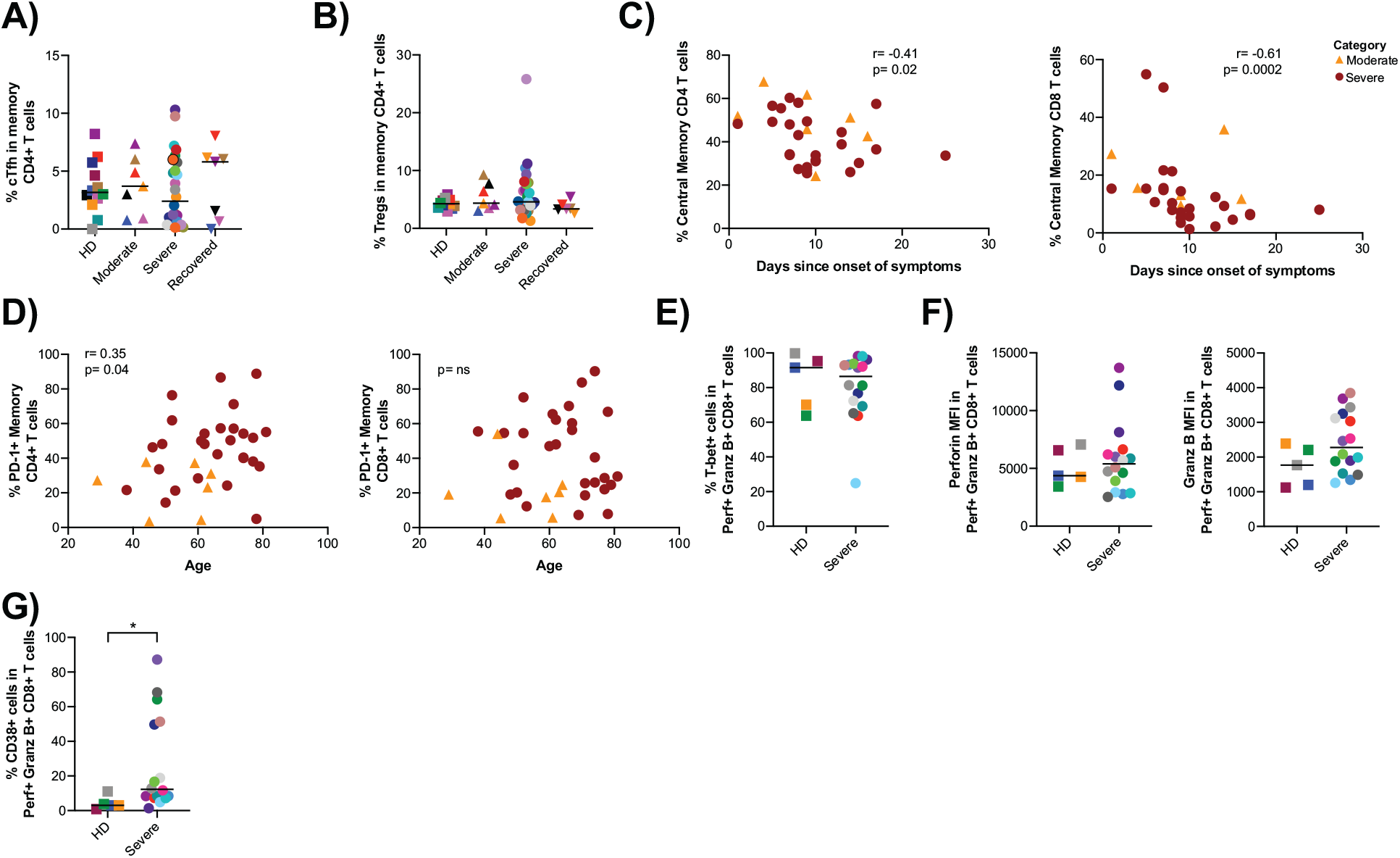
Extended T cell phenotype and activation during COVID-19 infection. **a-d)** Multiparametric flow cytometry analyses on fresh whole blood after red blood cell lysis characterizing immune cells subsets in HD (n= 12), moderate (n=7), severe (n=27), and recovered (n=6) COVID-19+ individuals was performed. Frequency of cTfh **(A)** and Tregs **(B)** (as defined in Fig. S1C). **C**) Spearman correlations of the frequency of CD4+ and CD8+ T_CM_ cells and days since onset of symptoms in moderate (orange triangles) and severe COVID-19+ individuals (dark red circles). **D)** Spearman correlations of the percentages of PD1+ CD4+ and CD8+ memory T cells and age in moderate and severe COVID-19+ individuals. **E-G)** Multiparametric flow cytometry analyses was performed on freshly isolated PBMC from HD (n=5) and severe (n=16) COVID-19+ individuals. **E)** Frequency of T-bet+ cells within cytotoxic CD8+ T cells (defined as granzyme B+ perforin+ memory CD8+ T cells). **F)** Expression of perforin and granzyme B (Mean fluorescence intensity) in cytotoxic CD8+ T cells. **G)** Frequency of activated cytotoxic CD8+ T cells. Specific color coding was assigned per individual for cross comparison across graphs and Figs. Lines on the graphs indicate the median of the group. Differences between groups were calculated using Kruskal-Wallis test with Dunn’s multiple comparison post-test, or Mann-Whitney rank sum test. *p<0.05, ns, not significant.

**Fig. S4.**
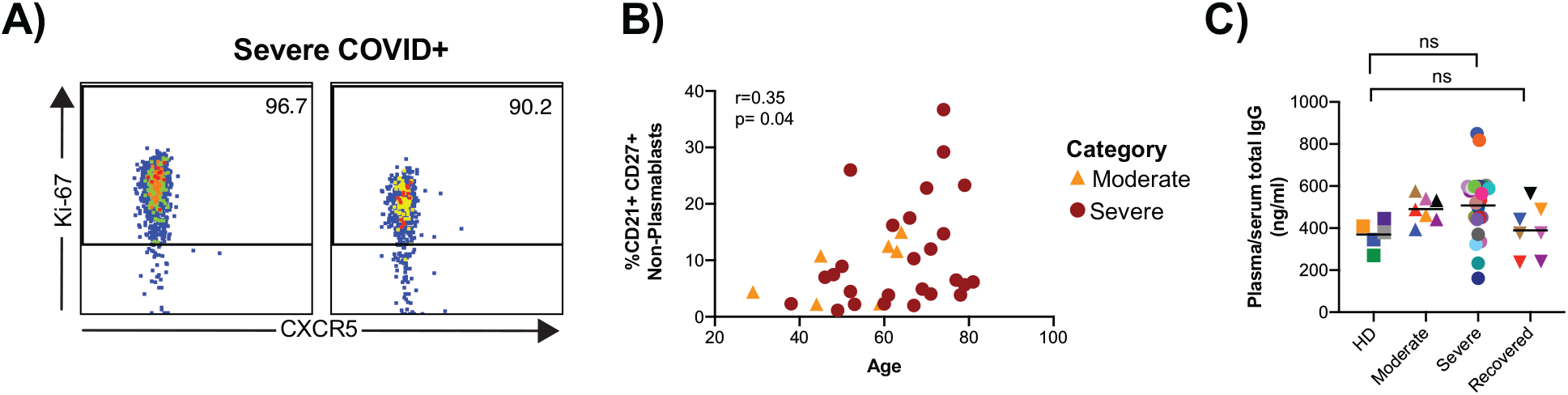
Extended B cell phenotype and total IgG measurements in COVID-19. **A)** Representative plots of the expression of Ki-67 and CXCR5 in plasmablasts in two severe COVID-19+ individuals. **B)** Spearman correlations of the frequency of CD21+CD27+ non-plasmablast B cells and age within moderate (orange triangles) and severe COVID-19+ individuals (dark red circles). **C)** Plasma/serum levels of total IgG measured in HD (n=5), moderate (n=7), severe (n=25) and recovered (n=7) COVID-19+ quantified using a cytometric bead array assay. Specific color coding was assigned per individual for cross comparison across graphs and Figs. Lines on the graphs indicate the median of the group. Differences between groups were calculated using Kruskal-Wallis test with Dun’s multiple comparison post-test. ns, not significant.

**Fig. S5.**
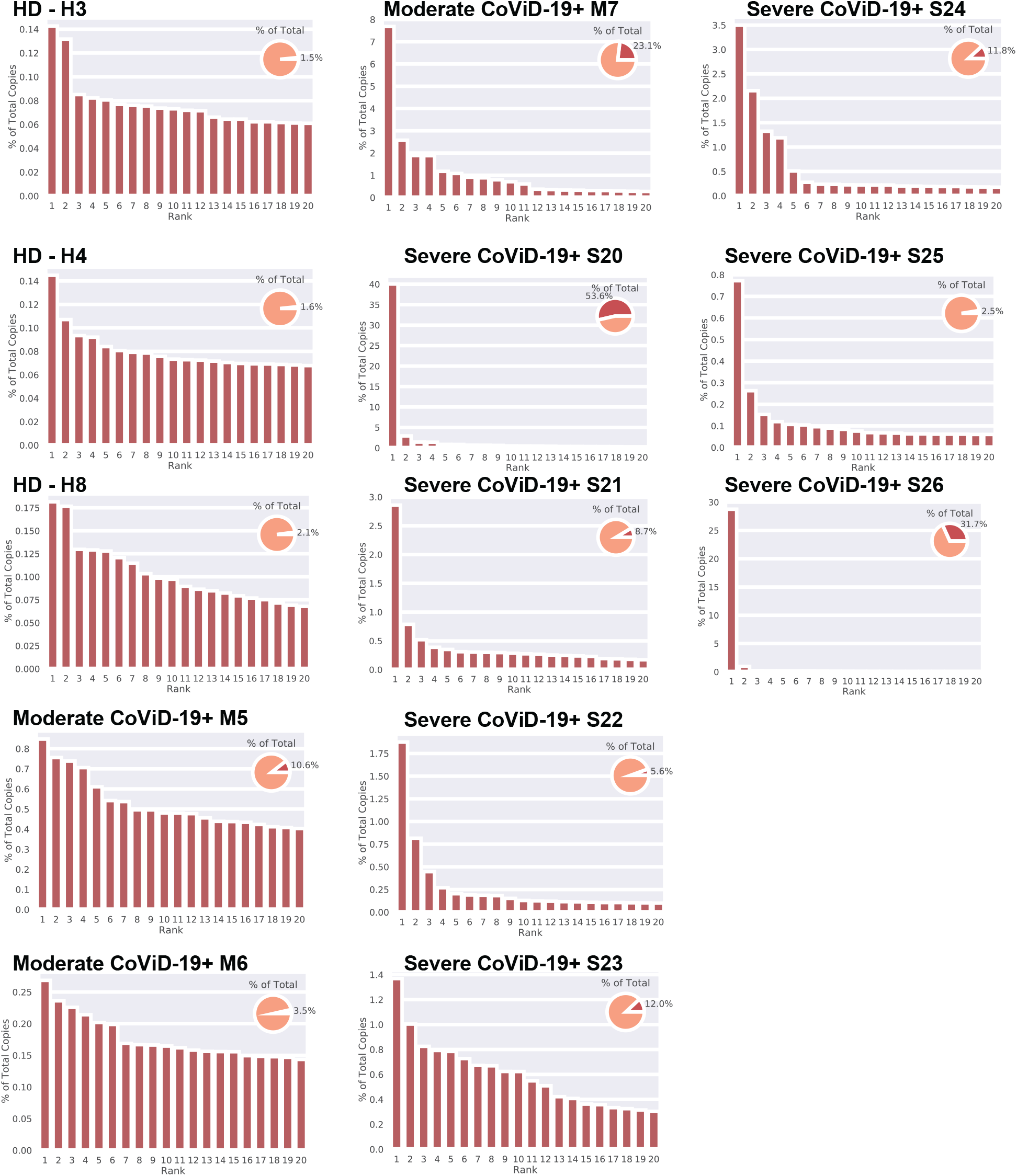
Abundance of the top 20 clones in each donor. The top twenty ranked clones and their copy number percentages are shown. Pie chart (inset) show the fraction of the total sequence copies that is comprised of the sum of the top 20 ranked clone copies (D20).

**Fig. S6.**
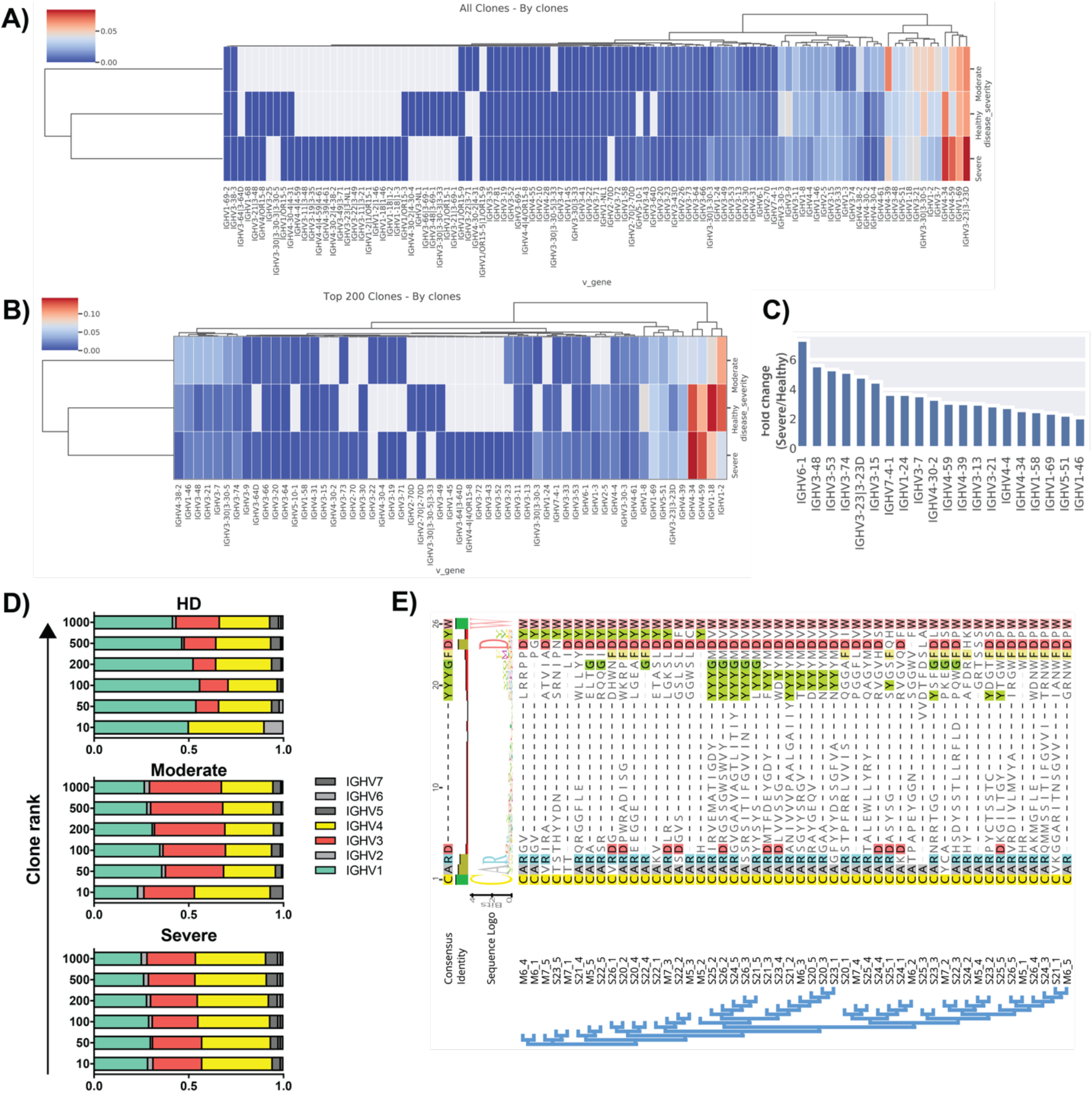
Heavy chain variable (VH) gene and CDR3 usage. **A)** VH usage of all clones, counting each clone only once per subject, data are aggregated and normalized by row (subject disease category); grey cells = no data. Data analyzed and visualized in ImmuneDB, see Materials and Methods. **B)** VH usage of the top 200 copy clones, counting each clone only once. **C)** Fold change in VH gene usage in COVID-19+ vs. HDs; analysis was limited to VH genes with at least one clone in both COVID-19+ and HDs; the top 20 VHs, ranked by fold change, are shown. **D)** VH family usage vs. binned clone ranks (10 = top ten copy number clones, 50 = top 50 copy number clones etc.) averaged over all individuals in each disease category. **E)** CDR3 amino acid alignment for top 5 copy clones in each of the severe and moderate COVID-19+ individuals, grouped by sequence similarity.

**Table S1.** Detailed clinical characteristics of individuals with moderate and severe COVID-19.

| Code | Donor | Cat | Days Sx Start | Age Bracket | Hypoxia Severity | APACHE III | Vasc/Metab Disorder<sup>a</sup> | Pulmonary Disorder<sup>b</sup> | Other infections/comorbidities |
| --- | --- | --- | --- | --- | --- | --- | --- | --- | --- |
| ▲ | M1 | Moderate | 4 | 61-65 | Room air | 42 | Y | N |  |
| ▲ | M2 | Moderate | 10 | 61-65 | NC | 34 | Y | Y | UTI gram positive |
| ▲ | M3 | Moderate | 4 | 56-60 | NC | 49 | Y | Y | Autoimmune disease on immunosuppression |
| ▲ | M4 | Moderate | 9 | 41-45 | Room air | 32 | Y | Y |  |
| ▲ | M5 | Moderate | 1 | 25-30 | Asymptomatic | 33 | N | N |  |
| ▲ | M6 | Moderate | 14 | 61-65 | Room air | 34 | Y | Y |  |
| ▲ | M7 | Moderate | 16 | 41-45 | NC | 20 | Y | N |  |
| ● | S1 | Severe | 10 | 46-50 | NIV / HFNC | 36 | Y | N |  |
| ● | S2 | Severe | 9 | 46-50 | Severe ARDS | 65 | Y | N |  |
| ● | S3 | Severe | 9 | 36-40 | Severe ARDS | 32 | Y | N | Pneumonia gram negative |
| ● | S4 | Severe | 17 | 51-55 | Severe-Mod ARDS | 50 | Y | N | Pneumonia gram positive |
| ● | S5 | Severe | 9 | 71-75 | Moderate ARDS | 72 | Y | N | Bacteremia gram positive |
| ● | S6 | Severe | 10 | 51-55 | Moderate ARDS | 23 | Y | N |  |
| ● | S7 | Severe | 1 | 71-75 | Ventilated non-ARDS | 112 | Y | N | Pneumonia gram positive |
| ● | S8 | Severe | 8 | 66-70 | Mild ARDS | 93 | N | N |  |
| ● | S9 | Severe | 10 | 46-50 | NIV / HFNC | 46 | Y | N |  |
| ● | S10 | Severe | 8 | 46-50 | Moderate ARDS | 32 | Y | Y | Solid organ transplant |
| ● | S11 | Severe | 7 | 66-70 | Severe ARDS | 69 | Y | N | Autoimmune disease on immunosuppression |
| ● | S12 | Severe | 15 | 71-75 | Mild ARDS | 83 | Y | N |  |
| ● | S13 | Severe | 13 | 66-70 | Moderate ARDS | 136 | Y | Y | Bacteremia gram negative and gram positive, UTI gram positive |
| ● | S14 | Severe | 5 | 66-70 | Mild ARDS | 78 | Y | N |  |
| ● | S15 | Severe | 7 | 56-60 | Severe ARDS | 130 | Y | Y |  |
| ● | S16 | Severe | 7 | 76-80 | Moderate ARDS | 129 | Y | Y |  |
| ● | S17 | Severe | 9 | 66-70 | Severe ARDS | 88 | Y | N | Immunocompromised, pneumonia gram negative |
|  | S18<sup>c</sup> | Severe | 10 | 76-80 | NIV / HFNC | 82 | Y | N | B cell lymphoma s/p rituxan |
|  | S19<sup>c</sup> | Severe | 9 | 76-80 | Severe ARDS | 156 | Y | N | B cell lymphoma |
| ● | S20 | Severe | 5 | 71-75 | Ventilated non-ARDS | 65 | Y | Y | Autoimmune disease |
| ● | S21 | Severe | 13 | 71-75 | Severe ARDS | 64 | Y | N | Bacteremia gram negative |
| ● | S22 | Severe | 17 | 76-80 | Moderate ARDS | 80 | Y | N |  |
| ● | S23 | Severe | 7 | 81-85 | Moderate ARDS | 76 | Y | N |  |
| ● | S24 | Severe | 25 | 51-55 | Severe ARDS - ECMO | 100 | Y | N | B cell lymphoma |
| ● | S25 | Severe | 14 | 61-65 | Severe ARDS | 80 | Y | Y | Bacteremia gram positive |
| ● | S26 | Severe | 8 | 71-75 | NIV / HFNC | 67 | Y | N |  |
| ● | S27 | Severe | 6 | 76-80 | Moderate ARDS | 66 | Y | Y |  |
| ● | S28 | Severe | 8 | 61-65 | Moderate ARDS | 76 | Y | N | Bacteremia gram positive |

Days Sx Start, days since onset of symptoms accounted from the time of blood draw. Hypoxia Severity: NC, nasal cannula; NIV / HFNC, non-invasive ventilation and/or high flow nasal cannula; ARDS, acute respiratory distress syndrome; ECMO, extracorporeal membrane oxygenation. APACHE, acute physiology and chronic health evaluation. One patient required mechanical ventilation for encephalopathy but did not fulfill ARDS radiographic criteria (“Ventilated non-ARDS”).
<sup>a</sup>Vascular and metabolic disorder category included any of the following: obesity, cardiovascular disease, hypertension, diabetes mellitus and hyperlipidemia. <sup>b</sup>Underlying pulmonary disorder category included asthma, sarcoidosis, chronic obstructive pulmonary disease or interstitial lung disease. Y, yes. N, no. <sup>c</sup>Excluded from all reported analyses.

**Table S2.**
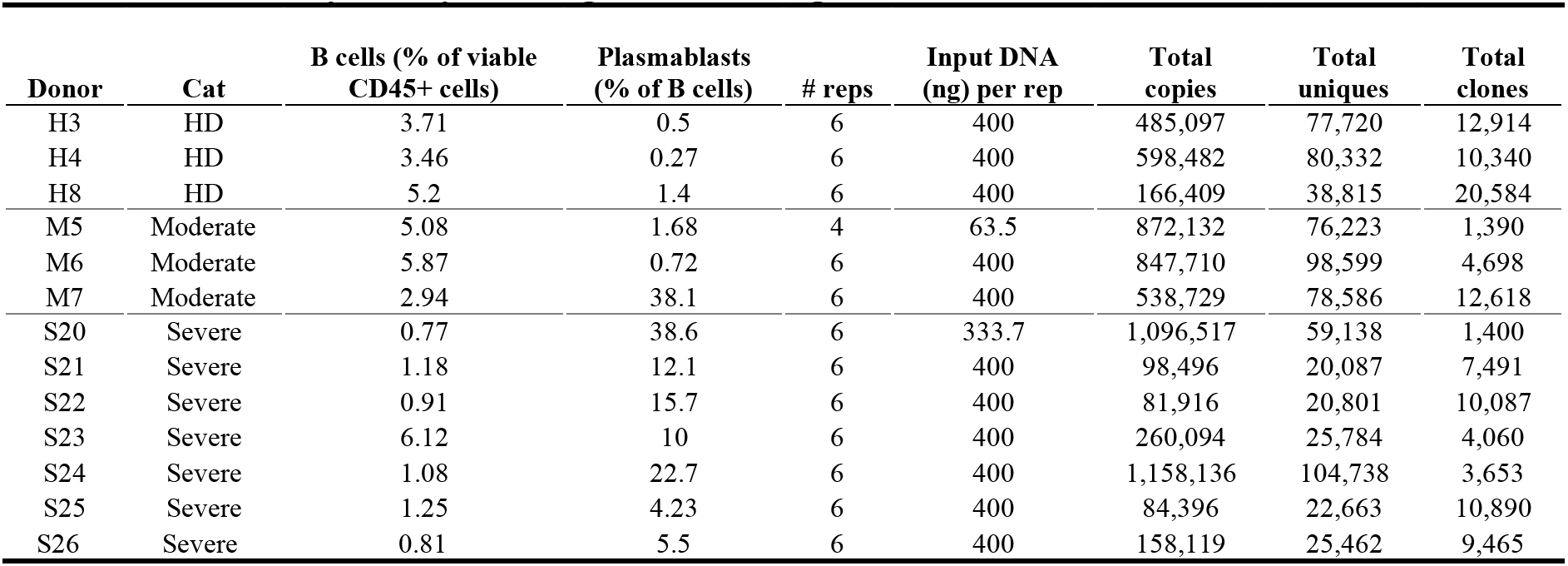
Antibody heavy chain gene rearrangement metadata. Frequencies of B cells and plasmablasts as characterized in Figure 2. Cat = disease category. HD = healthy donor. # reps, number of replicate sequencing libraries (independently amplified from genomic DNA). Total copies are sequence copies aggregated at the subject level. Total unique, unique sequences with each unique sequence variant counted only once across all replicate libraries from the same individual. Total clones, aggregated at the individual level with each clone only counted once. Clonally related sequences have the same VH gene and JH gene assignment, have identical third complementarity determining region (CDR3) sequence length and share 85% or more identity at the amino acid sequence level in the CDR3 (see Materials and Methods).

**Table S3.** Rotation table extracted from PCA. Top 20 elements extracted are shown for PC1 and PC2. All data are shown in percentages. T cell subsets defined as shown in Fig. S1B-C.

|  | PC1 | PC2 |
| --- | --- | --- |
| HLA-DR+ in TTM CD8+ | 0.163185775 | -0.03439971 |
| HLA-DR+ in TTM CD4+ | 0.153698234 | 0.001753146 |
| HLA-DR+ CD38+ in TCM CD4+ | 0.153608119 | 0.086296869 |
| HLA-DR+ CD38+ in TEM CD4+ | 0.152523955 | 0.047389036 |
| PD-1+ in TCM CD4+ | 0.150676319 | -0.024626777 |
| Plasmablasts in B cells | 0.145231599 | 0.063345638 |
| PD-1+ in TTM CD4+ | 0.144966645 | -0.067553736 |
| HLA-DR+ in TCM CD4+ | 0.144900513 | 0.081231632 |
| HLA-DR+ CD38+ in TTM CD4+ | 0.144418452 | -0.051306861 |
| PD-1+ in TTM CD8+ | 0.143632409 | -0.052141263 |
| PD-1+ in TEM CD4+ | 0.141261588 | -0.060674118 |
| CD38+ in TEM CD4+ | 0.140716008 | 0.044735059 |
| HLA-DR+ in TCM CD8+ | 0.13902669 | -0.048164109 |
| HLA-DR+ CD38+ in TTM CD8+ | 0.13822557 | -0.052535396 |
| Neutrophils in viable CD45+ | 0.13756484 | 0.019090674 |
| HLA-DR+ in TEM CD4+ | 0.136801039 | -0.006844502 |
| HLA-DR+ CD38+ total memory CD4+ | 0.135969253 | 0.036391482 |
| HLA-DR+ CD38+ total memory CD8+ | 0.134257028 | 0.127221692 |
| HLA-DR+ in TEMRA CD8+ | 0.130044138 | -0.06180719 |
| CD38+ in TTM CD8+ | 0.130034674 | -0.054027962 |

## References

1. W. Novel-Coronavirus-2019 Reports. (World Health Organization, 2020), vol. 2020.

2. W. J. Guan et al., Clinical Characteristics of Coronavirus Disease 2019 in China. N Engl J Med, (2020).

3. C. Huang et al., Clinical features of patients infected with 2019 novel coronavirus in Wuhan, China. Lancet 395, 497–506 (2020).

4. M. M. Lamers et al., SARS-CoV-2 productively infects human gut enterocytes. Science, eabc1669 (2020).

5. Z. Varga et al., Endothelial cell infection and endotheliitis in COVID-19. Lancet 395, 1417–1418 (2020).

6. X. H. Yao et al., [A pathological report of three COVID-19 cases by minimally invasive autopsies]. Zhonghua Bing Li Xue Za Zhi 49, E009 (2020).

7. N. Chen et al., Epidemiological and clinical characteristics of 99 cases of 2019 novel coronavirus pneumonia in Wuhan, China: a descriptive study. Lancet 395, 507–513 (2020).

8. Q. Ruan, K. Yang, W. Wang, L. Jiang, J. Song, Clinical predictors of mortality due to COVID-19 based on an analysis of data of 150 patients from Wuhan, China. Intensive Care Med, (2020).

9. X. Yang et al., Clinical course and outcomes of critically ill patients with SARS-CoV-2 pneumonia in Wuhan, China: a single-centered, retrospective, observational study. Lancet Respir Med, (2020).

10. D. Wang et al., Clinical Characteristics of 138 Hospitalized Patients With 2019 Novel Coronavirus-Infected Pneumonia in Wuhan, China. JAMA, (2020).

11. R. He et al., The clinical course and its correlated immune status in COVID-19 pneumonia. J Clin Virol 127, 104361 (2020).

12. J. J. Zhang et al., Clinical characteristics of 140 patients infected with SARS-CoV-2 in Wuhan, China. Allergy, (2020).

13. G. Chen et al., Clinical and immunological features of severe and moderate coronavirus disease 2019. J Clin Invest, (2020).

14. R. Wolfel et al., Virological assessment of hospitalized patients with COVID-2019. Nature, (2020).

15. I. Thevarajan et al., Breadth of concomitant immune responses prior to patient recovery: a case report of non-severe COVID-19. Nat Med 26, 453–455 (2020).

16. A. J. Wilk et al., A single-cell atlas of the peripheral immune response to severe COVID-19. medRxiv, 2020.2004.2017.20069930 (2020).

17. W. Wang et al., High-dimensional immune profiling by mass cytometry revealed immunosuppression and dysfunction of immunity in COVID-19 patients. Cellular & Molecular Immunology, (2020).

18. H.-Y. Zheng et al., Elevated exhaustion levels and reduced functional diversity of T cells in peripheral blood may predict severe progression in COVID-19 patients. Cellular & Molecular Immunology 17, 541–543 (2020).

19. D. R. Ziehr et al., Respiratory Pathophysiology of Mechanically Ventilated Patients with COVID-19: A Cohort Study. Am J Respir Crit Care Med, (2020).

20. A. D. T. Force et al., Acute respiratory distress syndrome: the Berlin Definition. JAMA 307, 2526–2533 (2012).

21. M. Merad, J. C. Martin, Pathological inflammation in patients with COVID-19: a key role for monocytes and macrophages. Nat Rev Immunol, (2020).

22. B. M. Henry, COVID-19, ECMO, and lymphopenia: a word of caution. Lancet Respir Med 8, e24 (2020).

23. L. Tan et al., Lymphopenia predicts disease severity of COVID-19: a descriptive and predictive study. Signal Transduct Target Ther 5, 33 (2020).

24. J. F. Chan et al., A familial cluster of pneumonia associated with the 2019 novel coronavirus indicating person-to-person transmission: a study of a family cluster. Lancet 395, 514–523 (2020).

25. J. Liu et al., Neutrophil-to-Lymphocyte Ratio Predicts Severe Illness Patients with 2019 Novel Coronavirus in the Early Stage. medRxiv, 2020.2002.2010.20021584 (2020).

26. B. Zhang et al., Immune phenotyping based on neutrophil-to-lymphocyte ratio and IgG predicts disease severity and outcome for patients with COVID-19. medRxiv, 2020.2003.2012.20035048 (2020).

27. D. Lau et al., Low CD21 expression defines a population of recent germinal center graduates primed for plasma cell differentiation. Sci Immunol 2, (2017).

28. J. Zhao et al., Antibody responses to SARS-CoV-2 in patients of novel coronavirus disease 2019. Clin Infect Dis, (2020).

29. G. E. Marti et al., Diagnostic criteria for monoclonal B-cell lymphocytosis. Br J Haematol 130, 325–332 (2005).

30. H. Tabibian-Keissar et al., Aging affects B-cell antigen receptor repertoire diversity in primary and secondary lymphoid tissues. Eur J Immunol 46, 480–492 (2016).

31. W. S. DeWitt et al., A Public Database of Memory and Naive B-Cell Receptor Sequences. PLoS One 11, e0160853 (2016).

32. A. Kurioka et al., CD161 Defines a Functionally Distinct Subset of Pro-Inflammatory Natural Killer Cells. Front Immunol 9, 486 (2018).

33. A. Poggi, A. Rubartelli, L. Moretta, M. R. Zocchi, Expression and function of NKRP1A molecule on human monocytes and dendritic cells. Eur J Immunol 27, 2965–2970 (1997).

34. C. W. Pohlmeyer et al., Identification of NK Cell Subpopulations That Differentiate HIV-Infected Subject Cohorts with Diverse Levels of Virus Control. J Virol 93, (2019).

35. M. R. Goodier et al., Sustained Immune Complex-Mediated Reduction in CD16 Expression after Vaccination Regulates NK Cell Function. Frontiers in Immunology 7, (2016).

36. E. J. Giamarellos-Bourboulis et al., Complex Immune Dysregulation in COVID-19 Patients with Severe Respiratory Failure. Cell Host Microbe, (2020).

37. S. Hua et al., Potential role for HIV-specific CD38-/HLA-DR+ CD8+ T cells in viral suppression and cytotoxicity in HIV controllers. PLoS One 9, e101920 (2014).

38. A. K. McElroy et al., Human Ebola virus infection results in substantial immune activation. Proc Natl Acad Sci U S A 112, 4719–4724 (2015).

39. Z. Wang et al., Clonally diverse CD38(+)HLA-DR(+)CD8(+) T cells persist during fatal H7N9 disease. Nat Commun 9, 824 (2018).

40. Z. Xu et al., Pathological findings of COVID-19 associated with acute respiratory distress syndrome. Lancet Respir Med 8, 420–422 (2020).

41. J. Braun et al., Presence of SARS-CoV-2 reactive T cells in COVID-19 patients and healthy donors. medRxiv, 2020.2004.2017.20061440 (2020).

42. A. R. Victor et al., Epigenetic and Posttranscriptional Regulation of CD16 Expression during Human NK Cell Development. J Immunol 200, 565–572 (2018).

43. K. Srpan et al., Shedding of CD16 disassembles the NK cell immune synapse and boosts serial engagement of target cells. J Cell Biol 217, 3267–3283 (2018).

44. J. M. Debets, C. J. Van der Linden, I. E. Dieteren, J. F. Leeuwenberg, W. A. Buurman, Fc-receptor cross-linking induces rapid secretion of tumor necrosis factor (cachectin) by human peripheral blood monocytes. J Immunol 141, 1197–1201 (1988).

45. Y. Liu et al., Phenotypic and clinical characterization of low density neutrophils in patients with advanced lung adenocarcinoma. Oncotarget 8, 90969–90978 (2017).

46. F. Nakayama et al., CD15 expression in mature granulocytes is determined by alpha 1,3-fucosyltransferase IX, but in promyelocytes and monocytes by alpha 1,3-fucosyltransferase IV. J Biol Chem 276, 16100–16106 (2001).

47. P. Bost et al., Host-viral infection maps reveal signatures of severe COVID-19 patients. Cell, (2020).

48. T. A. Mare et al., The diagnostic and prognostic significance of monitoring blood levels of immature neutrophils in patients with systemic inflammation. Crit Care 19, 57 (2015).

49. A. Nierhaus et al., Revisiting the white blood cell count: immature granulocytes count as a diagnostic marker to discriminate between SIRS and sepsis--a prospective, observational study. BMC Immunol 14, 8 (2013).

50. M. Liao et al., The landscape of lung bronchoalveolar immune cells in COVID-19 revealed by single-cell RNA sequencing. medRxiv, 2020.2002.2023.20026690 (2020).

51. T. Balakrishnan et al., Dengue virus activates polyreactive, natural IgG B cells after primary and secondary infection. PLoS One 6, e29430 (2011).

52. T. Dorner, P. E. Lipsky, Correlation of circulating CD27high plasma cells and disease activity in systemic lupus erythematosus. Lupus 13, 283–289 (2004).

53. M. Garcia et al., Massive plasmablast response elicited in the acute phase of hantavirus pulmonary syndrome. Immunology 151, 122–135 (2017).

54. J. Wrammert et al., Rapid and massive virus-specific plasmablast responses during acute dengue virus infection in humans. J Virol 86, 2911–2918 (2012).

55. M. Woodruff et al., Critically ill SARS-CoV-2 patients display lupus-like hallmarks of extrafollicular B cell activation. medRxiv, 2020.2004.2029.20083717 (2020).

56. H. Wardemann et al., Predominant autoantibody production by early human B cell precursors. Science 301, 1374–1377 (2003).

57. W. Wen et al., Immune Cell Profiling of COVID-19 Patients in the Recovery Stage by Single-Cell Sequencing. medRxiv, 2020.2003.2023.20039362 (2020).

58. S. M. Kahan, E. J. Wherry, A. J. Zajac, T cell exhaustion during persistent viral infections. Virology 479–480, 180–193 (2015).

59. C. Fenwick et al., T-cell exhaustion in HIV infection. Immunol Rev 292, 149–163 (2019).

60. C. N. S. Dias et al., Human CD8 T-cell activation in acute and chronic chikungunya infection. Immunology 155, 499–504 (2018).

61. Z. M. Ndhlovu et al., Magnitude and Kinetics of CD8+ T Cell Activation during Hyperacute HIV Infection Impact Viral Set Point. Immunity 43, 591–604 (2015).

62. K. R. Demers et al., Temporal Dynamics of CD8+ T Cell Effector Responses during Primary HIV Infection. PLoS Pathog 12, e1005805 (2016).

63. C. Agrati et al., Longitudinal characterization of dysfunctional T cell-activation during human acute Ebola infection. Cell Death Dis 7, e2164 (2016).

64. E. Sandalova et al., Contribution of herpesvirus specific CD8 T cells to anti-viral T cell response in humans. PLoS Pathog 6, e1001051 (2010).

65. X. Wang et al., SARS-CoV-2 infects T lymphocytes through its spike protein-mediated membrane fusion. Cell Mol Immunol, (2020).

66. P. Sarzi-Puttini et al., COVID-19, cytokines and immunosuppression: what can we learn from severe acute respiratory syndrome? Clin Exp Rheumatol 38, 337–342 (2020).

67. Y. X. Tan et al., Induction of apoptosis by the severe acute respiratory syndrome coronavirus 7a protein is dependent on its interaction with the Bcl-XL protein. J Virol 81, 6346–6355 (2007).

68. Y. Yue et al., SARS-Coronavirus Open Reading Frame-3a drives multimodal necrotic cell death. Cell Death Dis 9, 904 (2018).

69. L. M. Barton, E. J. Duval, E. Stroberg, S. Ghosh, S. Mukhopadhyay, COVID-19 Autopsies, Oklahoma, USA. Am J Clin Pathol 153, 725–733 (2020).

70. L. J. Walker et al., Human MAIT and CD8alphaalpha cells develop from a pool of type-17 precommitted CD8+ T cells. Blood 119, 422–433 (2012).

71. E. Billerbeck et al., Analysis of CD161 expression on human CD8^+^ T cells defines a distinct functional subset with tissue-homing properties. Proceedings of the National Academy of Sciences 107, 3006–3011 (2010).

72. B. van Wilgenburg et al., MAIT cells are activated during human viral infections. Nature Communications 7, 11653 (2016).

73. Y. Jouan et al., Functional alteration of innate T cells in critically ill Covid-19 patients. medRxiv, 2020.2005.2003.20089300 (2020).

74. E. E. Davenport et al., Genomic landscape of the individual host response and outcomes in sepsis: a prospective cohort study. Lancet Respir Med 4, 259–271 (2016).

75. V. Faivre, A. C. Lukaszewicz, D. Payen, Downregulation of Blood Monocyte HLA-DR in ICU Patients Is Also Present in Bone Marrow Cells. PLoS One 11, e0164489 (2016).

76. E. Guerin et al., Circulating immature granulocytes with T-cell killing functions predict sepsis deterioration*. Crit Care Med 42, 2007–2018 (2014).

77. I. N. Shalova et al., CD16 regulates TRIF-dependent TLR4 response in human monocytes and their subsets. J Immunol 188, 3584–3593 (2012).

78. G. Monneret et al., Persisting low monocyte human leukocyte antigen-DR expression predicts mortality in septic shock. Intensive Care Medicine 32, 1175–1183 (2006).

## References

1. A. D. T. Force et al., Acute respiratory distress syndrome: the Berlin Definition. JAMA 307, 2526–2533 (2012).

2. A. R. G. Flannery, S., Dhudasia, M.B, SARS-CoV-2 Seroprevalence Among Parturient Women. Research Square, (2020).

3. F. Amanat et al., A serological assay to detect SARS-CoV-2 seroconversion in humans. medRxiv, 2020.2003.2017.20037713 (2020).

4. D. Stadlbauer et al., SARS-CoV-2 Seroconversion in Humans: A Detailed Protocol for a Serological Assay, Antigen Production, and Test Setup. Curr Protoc Microbiol 57, e100 (2020).

5. W. Meng et al., An atlas of B-cell clonal distribution in the human body. Nat Biotechnol 35, 879–884 (2017).

6. J. A. Vander Heiden et al., pRESTO: a toolkit for processing high-throughput sequencing raw reads of lymphocyte receptor repertoires. Bioinformatics 30, 1930–1932 (2014).

7. A. M. Rosenfeld et al., Computational Evaluation of B-Cell Clone Sizes in Bulk Populations. Front Immunol 9, 1472 (2018).

8. J. Ye, N. Ma, T. L. Madden, J. M. Ostell, IgBLAST: an immunoglobulin variable domain sequence analysis tool. Nucleic Acids Res 41, W34–40 (2013).

9. A. M. Rosenfeld, W. Meng, E. T. Luning Prak, U. Hershberg, ImmuneDB, a Novel Tool for the Analysis, Storage, and Dissemination of Immune Repertoire Sequencing Data. Front Immunol 9, 2107 (2018).

10. D. A. Bolotin et al., MiXCR: software for comprehensive adaptive immunity profiling. Nat Methods 12, 380–381 (2015).

11. V. I. Levenshtein, Binary codes capable of correcting deletions, insertions and reversals. Soviet Physics Doklady 10, 707–710 (1966).

